# Meis1 isoform diversity orchestrates neural progenitor differentiation by regulating ATOH1 degradation at distinct subcellular compartments

**DOI:** 10.1101/2022.08.17.504235

**Authors:** Tomoo Owa, Toma Adachi, Ryo Shiraishi, Kentaro Ichijo, Kaiyuan Ji, Minami Mizuno, Kyoka Suyama, Kayo Nishitani, Ikuko Hasegawa, Masaki Sone, Daisuke Kawauchi, Tomoki Nishioka, Shinichiro Taya, Yutaka Suzuki, Kozo Kaibuchi, Satoshi Miyashita, Mikio Hoshino

## Abstract

The development of the complex nervous system is strictly controlled by diverse isoforms produced from individual genes, but the underlying machinery remains unclear. Our long-read cDNA sequencing identifies more than 700 genes with high isoform diversity in cerebellar granule cell progenitors (GCPs). One such gene, *Meis1*, produces MEIS1-FL and MEIS1-HdL isoforms, which include and lack the homeodomain, respectively. Our previous study showed that MEIS1-FL localizes to nuclei and promotes ATOH1 protein degradation through transcriptional regulation, thereby promoting GCP differentiation. In contrast, our *in vivo* electroporation experiment in this study shows that MEIS1-HdL inhibits GCP differentiation. MEIS1-HdL localizes in the cytoplasm and inhibits the degradation of ATOH1 mediated by CUL3, which is a newly identified E3 ligase for ATOH1. MEIS1-HdL enhances the binding of the COP9 signalosome to CUL3, which suppresses ATOH1 polyubiquitination. This study demonstrates that functionally antagonistic isoforms derived from a single gene cleverly control neural progenitor differentiation.

## Introduction

The complex and intricate central nervous system is constructed through many developmental steps. Its blueprint is thought to be largely encoded in genes, but the overall view of the genetic mechanisms of neural development remains largely unclear. Mammals, including humans, have only around 20,000 genes, but considering the number and types of neurons/glia and the complexity of neural network construction, this number may not be sufficient. Therefore, it is expected that multiple isoforms are produced from a single gene through alternative splicing and alternative transcription start sites, and play distinct roles. ^1,2^ Each isoform has a different molecular structure and is expressed at different developmental stages, in different cell types, or in different subcellular locations, enabling extremely complex control of neural development. ^3,4,5^ However, the isoforms registered in databases do not cover all actual isoforms, and it is believed that only a small fraction of isoforms have been identified so far. To capture all isoforms, conventional short-read RNA-sequencing (RNA-seq) is insufficient, and long-read RNA-seq is necessary. ^6,7^ However, long-read analyses have not been conducted much to date.

The development of the nervous system involves multiple stages, including the proliferation and differentiation of neural progenitors, neuronal migration, axon outgrowth and pathway exploration, and synapse formation, with many genes involved in each stage. To address the first step, the machinery for the proliferation/differentiation of neural progenitors, we investigated granule cell progenitors (GCPs) and granule cells (GCs) in the cerebellar development. During cerebellar development, GCPs located in the outer external granular layer (oEGL) initially express the transcription factor ATOH1 which maintain a more undifferentiated and proliferative state. ^8^ However, the ATOH1 protein is subject to precise degradation control, and as development progresses, GCPs lose ATOH1 expression and begin to express NEUROD1, becoming slightly differentiated with reduced proliferative capacity (ATOH1-nonexpressing GCPs). Although these cells were referred to as NEUROD1-positive GCPs, we call them ATOH1-nonexpressing GCPs in this study. ATOH1-nonexpressing GCPs subsequently become postmitotic and differentiate into GCs. ^9^ This transition from ATOH1-expressing GCPs to ATOH1-nonexpressing GCPs is strictly controlled by the degradation of the ATOH1 protein, but the mechanism controlling ATOH1 degradation has not yet been fully elucidated. ^8,10,11^

In this study, we perform long-read RNA-seq on cerebellar GCPs to reveal extensive isoform diversity, identifying more than 700 genes with multiple isoform variants. Among genes showing high diversity, we have selected *Meis1* as a model gene, investigating its two distinct isoforms. One is the full-length isoform (MEIS1-FL), which has been studied extensively as a transcription factor with a homeodomain, and the other is a functionally unknown isoform (MEIS1-HdL) that lacks a homeodomain. MEIS1 (MEIS1-FL) is a homeodomain transcription factor belonging to the TALE family and is known to play an important role in stem cell maintenance, organ formation, and cell differentiation in various developmental processes. ^12,13,14,15^ Our previous gene-level analysis using the loss-of-function mutant mice for *Meis1* showed that MEIS1-FL activates *Pax6* transcription, enhances the BMP signaling, and promotes ATOH1 degradation, thereby enhancing the differentiation of GCPs into GCs. ^16^ However, these investigations did not distinguish between the functions of its specific isoforms, leaving their individual roles unexplored. The two isoforms, in fact, show striking differences in their expression patterns and subcellular localization: MEIS1-FL is expressed in both GCPs and GCs, and is localized in the cell nuclei, whereas MEIS1-HdL is expressed in GCPs but not in GCs, and is localized in the cytoplasm. Therefore, we hypothesized that MEIS1-HdL might have significantly different functions from MEIS1-FL in GCP proliferation and differentiation.

Here, we investigate the roles of these MEIS1 isoforms in the regulation of GCP proliferation and differentiation, with a particular focus on the degradation control of ATOH1. Our *in vivo* experiments show that the two isoforms play opposite roles in precise regulation of GCP proliferation and differentiation. Each isoform functions in a distinct intracellular location and thereby exerts an opposite effect on ATOH1 degradation. This study provides an example of how multiple isoforms derived from a single gene precisely control neurogenesis and contributes to our further understanding of the molecular mechanisms of neurogenesis, as well as the degradation control mechanisms of critical proteins.

## Results

### Comprehensive long-read cDNA sequencing of cerebellar granule cell progenitors reveals extensive isoform diversity

To comprehensively investigate the isoform landscape in cerebellar GCPs, we performed long-read cDNA sequencing on GCP samples using a Nanopore sequencer ^17^ (Figure 1A). Our analysis identified a total of 99,010 transcripts (Figure 1B). Among the identified transcripts, known isoforms accounted for a total of 58.3%, comprising 47.7% Full Splice Match (FSM) isoforms that perfectly matched existing annotations, and 10.6% Incomplete Splice Match (ISM) isoforms (Figure 1B). We also detected a substantial number of novel isoforms, with 14.8% being Novel In Catalog (NIC) and 17.8% being Novel Not In Catalog (NNIC) (Figure 1B). The remaining transcripts included categories such as Genic Genomic, Antisense, Fusion, Intergenic, and Genic Intron. This broad range of isoform categories highlights the extensive complexity of the GCP transcriptome captured by long-read sequencing.

**Figure 1.**
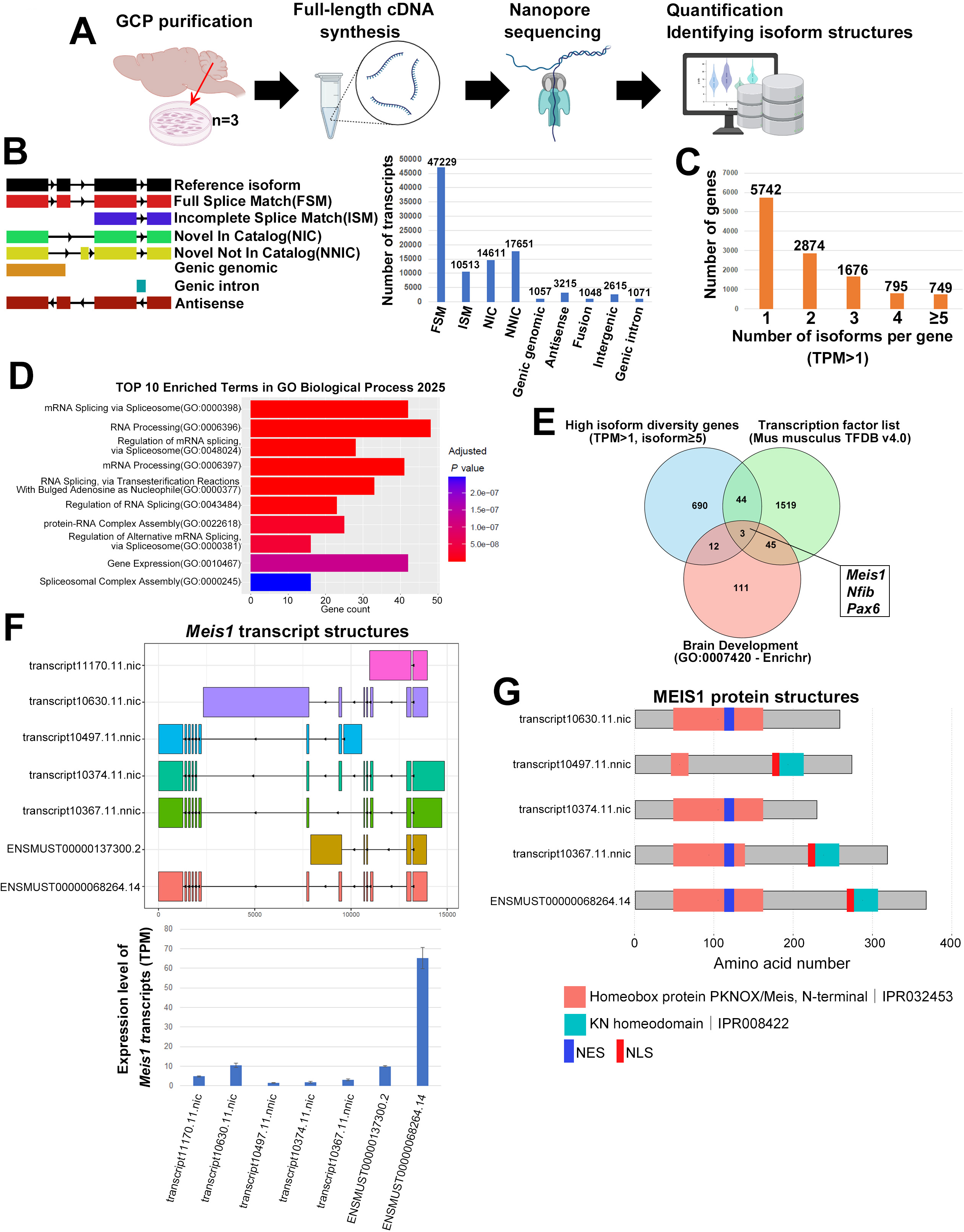
Comprehensive long-read cDNA sequencing unveils extensive isoform diversity and key transcription factors in cerebellar GCPs. **A.** Schematic diagram of the cDNA long-read sequencing analysis workflow using cerebellar granule cell progenitors (GCPs). **B.** Classification of all identified transcripts (99,010 in total) based on Sqanti3 ^58^ categories, displaying the proportion of Full Splice Match (FSM), Incomplete Splice Match (ISM), Novel In Catalog (NIC), Novel Not In Catalog (NNIC), and other categories. **C.** Distribution of isoform counts per gene for robustly expressed genes (TPM > 1 in all three replicates). The graph shows the percentage of genes expressing 1, 2, 3, 4, or ≥5 isoforms (11,836 genes in total). **D.** Gene Ontology (GO) enrichment analysis for genes with high isoform diversity (defined as ≥5 isoforms with TPM > 1 in all three replicates; 749 genes total), performed using Enrichr ^60,61,62^ (GO Biological Process 2025). **E.** Venn diagram showing the intersection of three gene lists to identify key transcription factors relevant to cerebellar development: (1) genes with high isoform diversity, (2) known transcription factors (from Mus musculus TFDB v4.0), and (3) genes associated with “Brain Development” term (GO:0007420) from the Enrichr. The analysis identified three common transcription factors: *Meis1*, *Pax6*, and *Nfib*. The complete list of high-diversity genes is provided in the Supplementary File. **F.** Structures and expression levels of identified *Meis1* transcript isoforms. Top: Isoform structures visualized with ggtranscript. ^63^ Bottom: Corresponding expression levels in Transcripts Per Million (TPM). **G.** Predicted protein domain structures of coding *Meis1* isoforms. Protein sequences were obtained via Sqanti3, with domains predicted by InterPro ^64^ and visualized using drawProteins. ^65^ Key conserved domains are shown. Locations of the putative Nuclear Export Signal (NES; blue) and Nuclear Localization Signal (NLS; red) were manually annotated. Some isoforms lack the C-terminal homeodomain containing the NLS.

To focus our analysis on robustly expressed transcripts, we next assessed their expression levels. A histogram of Transcript Per Million (TPM) values revealed that a substantial fraction of transcripts, particularly the novel isoforms, were expressed at low levels (Figure S1C). Since low-abundance transcripts can represent either rare, functional isoforms or transcriptional noise, we applied a stringent filter to create a high-confidence dataset for downstream analysis.

To understand the extent of isoform diversity per gene, we analyzed the number of isoforms expressed by each gene. Initially, without filtering by expression level, we found that a substantial proportion of genes (30.86%) expressed five or more isoforms (Figure S1D). To focus on robustly expressed isoforms, we applied a stringent filter, retaining only those transcripts with a TPM value greater than 1 in each of our three replicates. This analysis identified 11,836 robustly expressed genes. Of these, 48.5% expressed a single isoform, while a notable 6.33% of genes (749 genes) expressed five or more isoforms, highlighting a subset of genes with particularly high isoform diversity (Figure 1C).

We then performed Gene Ontology (GO) enrichment analysis for the 749 genes exhibiting high isoform diversity (≥5 isoforms with TPM > 1), focusing on Biological Process terms (Figure 1D). The top enriched GO terms were predominantly related to mRNA splicing (e.g., “mRNA Splicing via Spliceosome,” “RNA Processing,” “Regulation of mRNA splicing”) and protein–RNA complex assembly. This enrichment suggests that genes with high isoform diversity are themselves enriched in functions related to the regulation and execution of transcript diversification, indicating a self-reinforcing mechanism of complexity in the transcriptome.

### Identification of transcription factors with extensive isoform diversity in GCPs

To understand how this extensive isoform diversity regulates GCP differentiation, we focused on transcription factors, as they are the master regulators that orchestrate the gene expression programs driving this process. We therefore performed a Venn diagram analysis (Figure 1E). This analysis combined the 749 high isoform diversity genes with a list of transcription factors and 171 genes annotated with the “Brain Development” GO term (GO:0007420). This intersection revealed three common transcription factors: *Meis1*, *Pax6*, and *Nfib*. All of these transcription factors have been implicated in cerebellar development, with their disruption leading to structural or developmental abnormalities. ^16,18,19, 20^

For *Meis1*, we detected seven transcripts (five novel), predicted to encode several protein variants. Most notably, these included a canonical full-length isoform and a major novel isoform lacking the DNA-binding homeodomain (Figures 1F, 1G). ^21^ For *Pax6*, we identified ten transcripts (four novel). Our data captured its well-established diversity, including known isoforms like *Pax6(5a)* that arise from subtle variations within the paired domain to alter DNA-binding specificity. ^22,23^ In addition to these, our analysis revealed more drastically altered novel variants, including the one that lacks the paired domain entirely while retaining the homeodomain, and another predicted to be a short protein lacking both canonical domains (Figures S1E, F). For *Nfib*, we detected fourteen transcripts (eight novel). This aligns with the known strategy for *Nfib*, where functional diversity often arises from variations in the C-terminal transactivation domain while the N-terminal DNA-binding domain is conserved, generating proteins such as dominant-negative isoforms. ^24,25^ Our analysis expanded this repertoire, uncovering novel variants with more substantial changes, including one completely lacking the CTF/NFI C-terminal domain itself, and another lacking all predicted domains (Figures S1G, H). Among these three factors, we chose to focus on *Meis1* for our in-depth functional analysis. This decision was based on two key points. First, in contrast to the isoforms of *Pax6* and *Nfib* that primarily suggest changes in transcriptional activity, the diversity of *Meis1* isoforms pointed to a novel regulatory mechanism based on subcellular localization. Specifically, we found that a canonical full-length isoform containing a nuclear localization signal (NLS) and a major homeodomain-less variant that lacks the NLS but retains domains associated with a nuclear export signal (NES) were co-expressed in GCPs. ^12^ Second, *Meis1*’s established role as an upstream regulator of *Pax6* placed this potential mechanism in a critical biological context. Therefore, we proceeded to use *Meis1* as a model to understand how the structural diversity of its isoforms orchestrates the process of neural progenitor differentiation.

### *Meis1* gene produces two major, spatially separated protein isoforms in GCPs

To examine the protein expression of MEIS1 isoforms in GCPs, we performed Western blotting with a pan-MEIS1 antibody (Figure 2A, left panel). We observed a major band around 50 kDa, consistent with the predicted size of the full-length protein, and two distinct lower molecular weight bands at approximately 32 kDa and 27 kDa. As these were the only major protein products robustly detected, our subsequent functional analysis focused on these forms. This pattern of ∼27 kDa and ∼32 kDa bands was consistent with a previously described homeodomain-less isoform, “MEIS1D”, reported in colorectal cancer. ^21^ Based on this, we hypothesized that the lower molecular weight bands in GCPs represent unmodified and post-translationally modified forms of this homeodomain-less isoform. To test this, we generated a specific antibody against the unique 5-amino acid C-terminus of the predicted homeodomain-less isoform. After validating the specificity of our newly generated MEIS1 homeodomain-less specific antibody (Figure S2A), we verified that the ∼27 kDa and ∼32 kDa bands were indeed the homeodomain-less isoform in GCP lysate (Figure 2A, middle panel). Based on this protein evidence, we hereafter refer to the full-length protein as MEIS1-FL and the homeodomain-less isoform as MEIS1-HdL (*transcript10374.11.nic* in Figure 1G). Thus, Western blot and RT-PCR analyses collectively demonstrate that GCPs co-express two major Meis1 isoforms, FL and HdL, at both the protein and transcript levels (Figure 2A, B).

**Figure 2.**
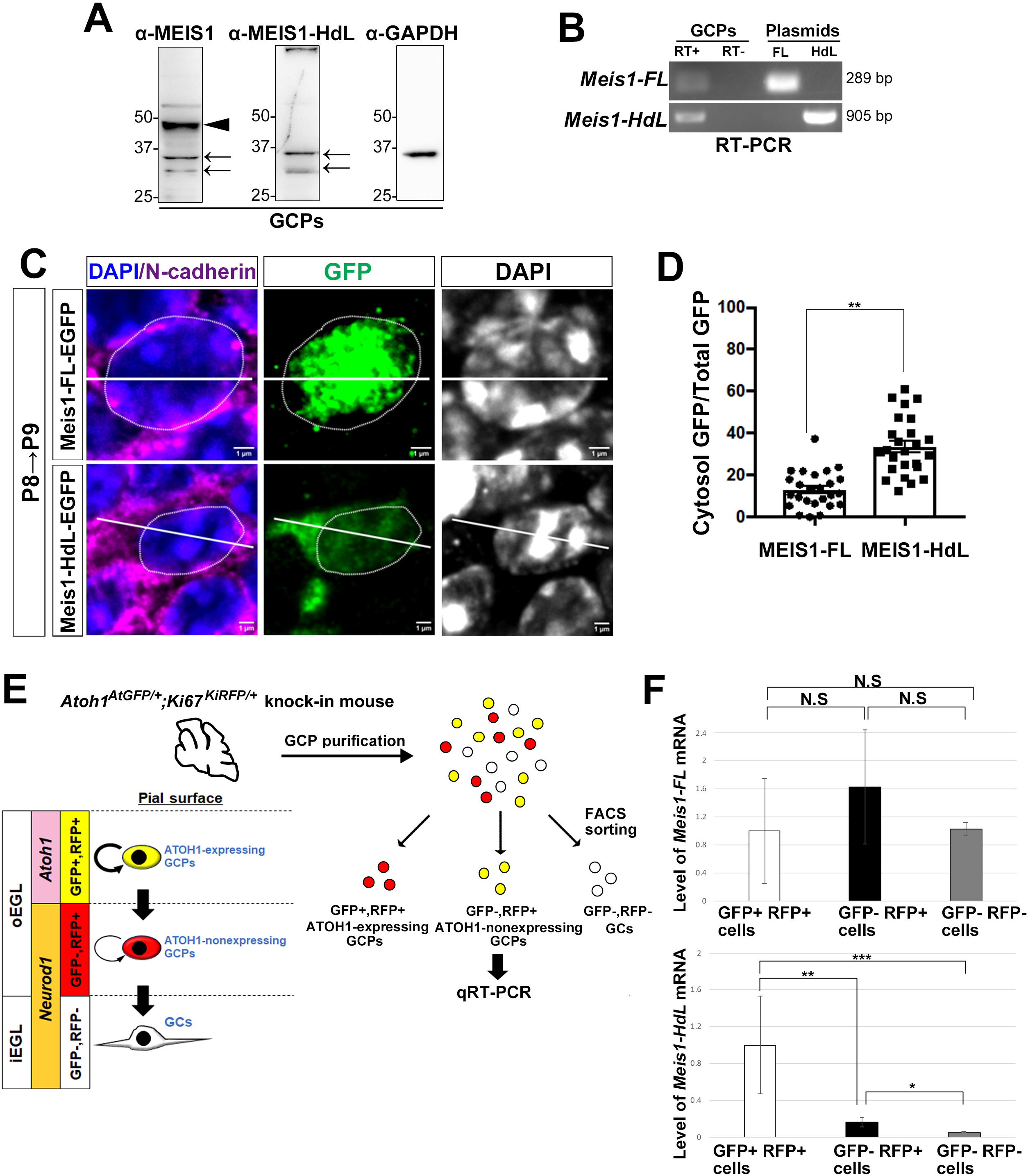
Meis1 isoforms exhibit distinct expression patterns and subcellular localizations in GCPs. **A.** Immunoblotting of P7 GCP cell lysates using the indicated antibodies. An arrowhead indicates the MEIS1-FL band (∼50 kDa), and arrows indicate two distinct bands for MEIS1-HdL (∼32 kDa and ∼27 kDa). **B.** *Meis1* PCR amplification using a negative control (RT^−^), plasmids coding for Meis1-FL or Meis1-HdL, or P6 GCP cDNA (RT^+^) templates. Upper panel: RT-PCR performed with a primer pair that amplifies only Meis1-FL (primers recognizing exon8 and exon11). Lower panel: RT-PCR was performed using primer pairs that amplify only Meis1-HdL (primers recognizing the start codon and the junction of exon7-9). Both Meis1 isoforms are detected by RT-PCR. **C.** Representative immunofluorescence images showing the subcellular localization of MEIS1-FL-EGFP and MEIS1-HdL-EGFP in P9 cerebellar GCPs electroporated *in vivo* at P8. Sections were immunostained for GFP (green, MEIS1 isoforms), N-cadherin (magenta, cell boundaries), and counterstained with DAPI (blue, nuclei). **D.** Quantification of the cytosolic-to-total GFP signal ratio in individual electroporated cells from (C). The cytosolic region was defined as the N-cadherin-positive cellular area excluding the DAPI-stained nucleus. **E.** Schematic of GCP separation via FACS, using *Atoh1^AtGFP^/^+^; Ki67^KiRFR^/^+^*mice. This method isolates three distinct populations used for subsequent analysis: ATOH1-expressing GCPs (GFP+RFP+), ATOH1-nonexpressing GCPs (GFP-RFP+), and postmitotic granule cells (GCs; GFP-RFP-). **F.** Relative mRNA expression of *Meis1-FL* and *Meis1-HdL* in the cell populations defined in (E), as quantified by qRT-PCR.

Next, we investigated the subcellular localization of MEIS1 isoforms in GCPs. Subcellular fractionation of GCP lysates followed by Western blotting with a pan-MEIS1 antibody revealed that the ∼50 kDa band (MEIS1-FL) was predominantly localized in the nuclear fraction, while the lower bands (MEIS1-HdL) were found primarily in the cytoplasmic fraction (Figure S2B). This observation aligns with our structural prediction that MEIS1-FL contains an NLS within its homeodomain, whereas MEIS1-HdL, lacking the homeodomain, retains an NES (Figure 1G). To investigate this differential localization in the developing cerebellum, we introduced plasmids encoding MEIS1-FL-EGFP and MEIS1-HdL-EGFP fusion proteins into GCPs in the postnatal day 8 (P8) cerebellum via *in vivo* electroporation, and analyzed them at P9. Immunofluorescence staining showed that MEIS1-FL-EGFP localized strongly to the nucleus, while MEIS1-HdL-EGFP exhibited robust cytoplasmic localization (Figure 2C, D). We obtained consistent results in N2a cells. Subcellular fractionation of N2a cells overexpressing untagged constructs revealed that MEIS1-FL was largely nuclear but also present in the cytoplasm, whereas MEIS1-HdL was predominantly cytoplasmic (Figure S2C). Consistent with this, immunofluorescence staining of N2a cells overexpressing EGFP-tagged constructs (Figure S2D) further confirmed that MEIS1-FL-EGFP localized strongly to the nucleus, while MEIS1-HdL-EGFP showed robust cytoplasmic localization. Taken together, these results demonstrate that MEIS1 isoforms occupy distinct subcellular compartments in GCPs, and this physical separation strongly suggests they have distinct functional roles.

### Differential expression profiles of *Meis1-FL* and *Meis1-HdL* during GCP differentiation

We next examined the developmental stage-specific expression of *Meis1-FL* and *Meis1-HdL* in GCPs. Our previous work has shown a two-step amplification model for GCP differentiation, where highly proliferative, ATOH1-expressing GCPs give rise to an intermediate, transit-amplifying population that is ATOH1-negative and NEUROD1-positive, before finally exiting the cell cycle to become GCs. ^9^ To distinguish and isolate these distinct progenitor populations, we utilized *Atoh1^AtGFP/+^; Ki67^KiRFP /+^* knock-in mice. ^26,27^ We validated these reporter mice by demonstrating that GFP signal largely overlapped with Atoh1 immunostaining (Figure S3A), and RFP signal overlapped with KI67 immunostaining (Figure S3B). The combined GFP and RFP expression patterns showed that GFP was more restricted to the oEGL, while RFP was expressed more broadly but still predominantly in the oEGL (Figure S3C). This finding is consistent with the known restricted localization of endogenous ATOH1 to the oEGL, ^8^ further validating our reporter mice. We used FACS to isolate three populations based on our model: ATOH1-expressing GCPs (GFP+RFP+), ATOH1-nonexpressing GCPs (GFP-RFP+), and postmitotic GCs (GFP-RFP-) (Figure 2E, S3D). The specificity of our sorting strategy was further validated by qRT-PCR for *Atoh1*, which showed *Atoh1* expression exclusively in GFP+RFP+ sorted cells (Figure S3E). Quantitative RT-PCR analysis of *Meis1* isoforms across these sorted populations revealed that *Meis1-HdL* expression was highly enriched in GFP+RFP+ sorted cells (ATOH1-expressing GCPs), with minimal expression in GFP-RFP+ sorted GCPs (ATOH1-nonexpressing, NEUROD1-expressing GCPs) and GFP-RFP-sorted cells (GCs) (Figure 2F, lower graph). In contrast, *Meis1-FL* was expressed at relatively constant levels across ATOH1-expressing GCPs, ATOH1-nonexpressing GCPs, and postmitotic GCs (Figure 2F, upper graph). Taken together with our localization data, these findings reveal a clear spatio-temporal partitioning of the two major MEIS1 protein products. MEIS1-HdL is confined to the cytoplasm and is predominantly expressed at the earliest, most proliferative stage of GCP development. This physical and developmental segregation suggests they have distinct functional roles.

### Opposing functions of MEIS1-FL and MEIS1-HdL in regulating GCP fate *in vivo*

To examine the functions of MEIS1-FL and MEIS1-HdL in cerebellar granule cell development, we performed *in vivo* overexpression experiments in GCPs. We introduced expression vectors encoding either Meis1-FL or Meis1-HdL into ATOH1-expressing GCPs via electroporation into the P8 cerebella. At 72 hours post-electroporation (P11), the cerebella were collected, fixed, and immunostained. Electroporated cells were identified by co-electroporated histone H3.1-GFP.

In control animals, the GFP+ cells comprised approximately 50% GCPs (Ki67+ cells) and 50% differentiating/differentiated GCs (p27+ cells) (Figure 3A, D, E and Figure S4A). Overexpression of Meis1-FL significantly decreased the proportion of GCPs (Ki67+ cells) and increased the proportion of GCs (p27+ cells) (Figure 3B, D, E and Figure S4B). Conversely, Meis1-HdL overexpression led to an increase in the proportion of GCPs (Ki67+ cells) and a decrease in the proportion of GCs (p27+ cells), indicative of maintaining an undifferentiated, proliferative state (Figure 3C, D, E and Figure S4C). Similarly, MEIS1-FL overexpression decreased the proportion of ATOH1-expressing GCPs, whereas MEIS1-HdL overexpression increased ATOH1-expressing GCPs (Figure 3A-C, F). These results indicate that MEIS1-FL and MEIS1-HdL have opposing effects on GCP differentiation. This promotion of GCP differentiation by MEIS1-FL is consistent with our previous observation that conditional deletion of *Meis1*, an allele that disrupts all homeodomain-containing isoforms (most notably the full-length MEIS1-FL), delayed GCP differentiation. ^16^

**Figure 3.**
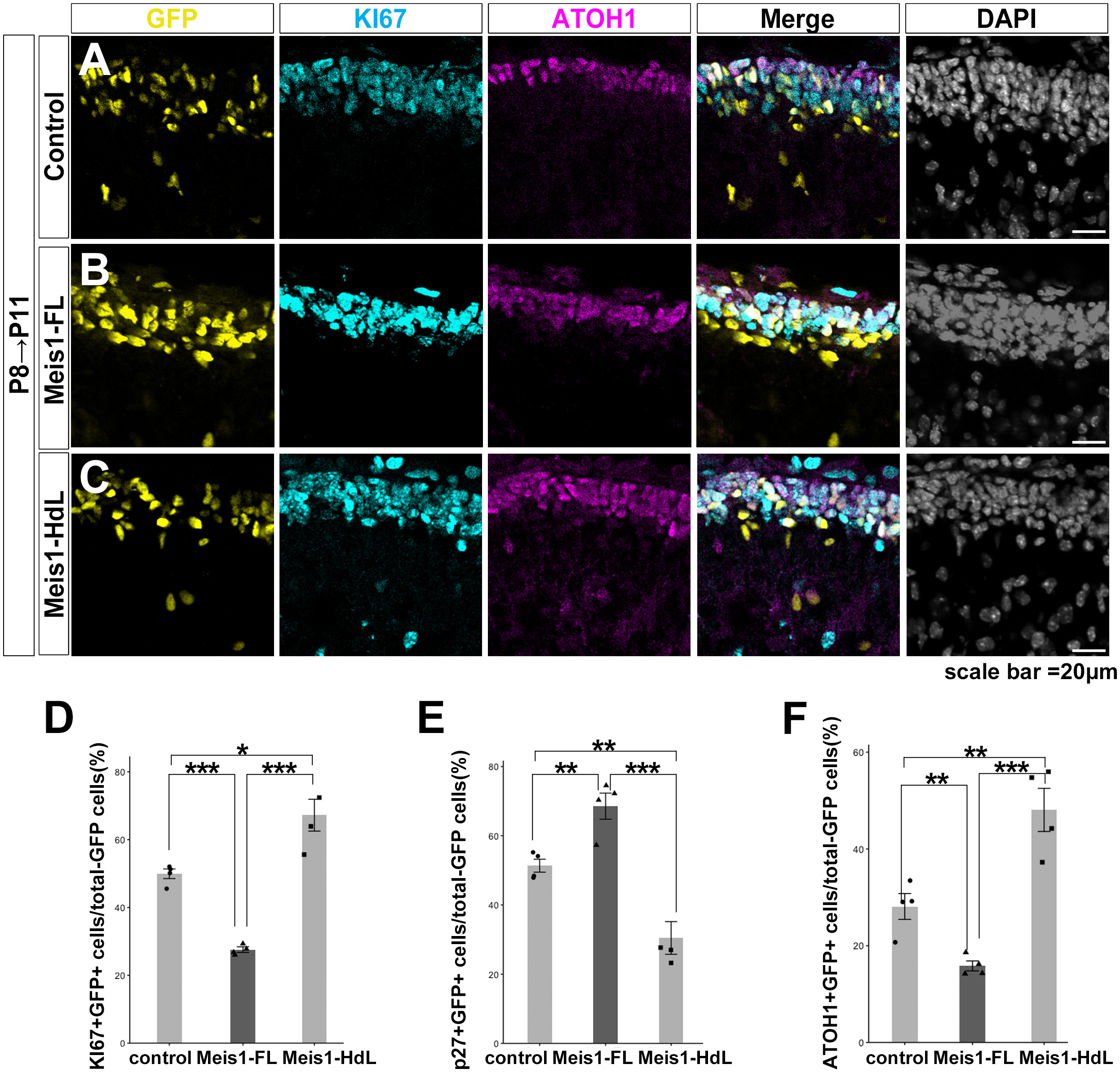
MEIS1-HdL maintains the undifferentiated state of GCPs, while MEIS1-FL promotes GC differentiation. **A–C.** Representative immunofluorescence images of P11 cerebella following *in vivo* electroporation at P8. Cerebellar sections were immunostained for ATOH1 (magenta) and KI67 (cyan). Electroporated cells, identified by co-electroporated H3.1-EGFP (yellow), were transfected with a control vector (A), Meis1-FL (B), or Meis1-HdL (C). **D–F.** Quantification of electroporated cell differentiation status. The percentage of GFP+ cells expressing KI67 (D), p27 (E), or ATOH1 (F) was quantified from sections described in (A–C).

To further investigate the physiological function of MEIS1-HdL, we performed loss-of-function experiments using shRNA. We first generated a *Meis1-HdL* specific knockdown vector (sh-*Meis1-HdL),* which effectively suppressed *Meis1-HdL* expression without affecting *Meis1-FL* (Figure S5A, left panel). For comparison, we also utilized a previously established *pan-Meis1* knockdown vector (sh-*Meis1-all*), ^16^ which suppressed the expression of both *Meis1-FL* and *Meis1-HdL* (Figure S5A, right panel).

These shRNA vectors, along with co-electroporated histone H3.1-GFP to identify electroporated cells, were introduced into GCPs in the P8 cerebella via *in vivo* electroporation. Cerebellar sections were analyzed by immunostaining 72 hours post-electroporation (P11). Specific knockdown of *Meis1-HdL* significantly reduced the proportion of proliferative GCPs (Ki67+ cells) and increased the proportion of postmitotic GCs (p27+ cells) (Figure 4A, D, E and Figure S5B). A similar trend was observed for ATOH1-expressing GCP population (Figure 4A, F). These findings suggest that the endogenous function of MEIS1-HdL is to suppress GCP differentiation. Crucially, the phenotypes induced by sh-*Meis1-HdL* were rescued by co-electroporation with a knockdown-resistant Meis1-HdL expression vector (Res-Meis1-HdL), restoring the proportion of KI67+ (Figure 4B, D) and ATOH1+ cells (Figure 4B, F), and decreasing p27+ cells (Figure S5C and Figure 4E). This rescue demonstrates that the observed effects are due to the loss of MEIS1-HdL, thereby establishing its critical role in suppressing GCP differentiation.

**Figure 4.**
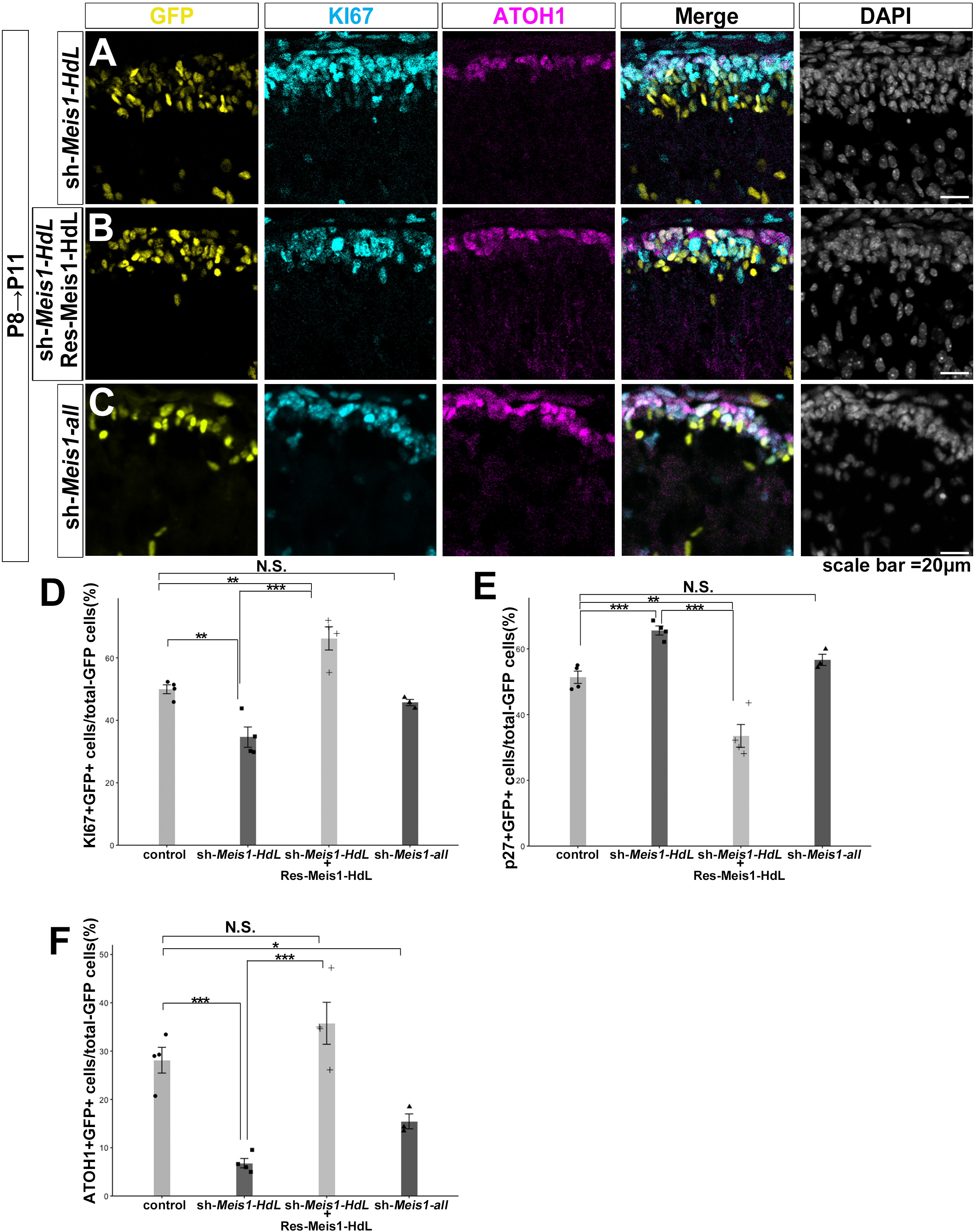
Endogenous MEIS1-HdL participates in maintaining GCPs in an immature and proliferative state. **A–C.** Representative immunofluorescence images of P11 cerebella following *in vivo* electroporation at P8. Cerebellar sections were immunostained for ATOH1 (magenta) and KI67 (cyan). Electroporated cells, identified by co-electroporated H3.1-EGFP (yellow), were transfected with sh-*Meis1-HdL* (A), sh-*Meis1-HdL* plus a knockdown-resistant rescue construct (Res-Meis1-HdL) (B), or sh-*Meis1-all* (C). **D–F.** Quantification of electroporated cell differentiation status. The percentage of GFP+ cells expressing KI67 (D), p27 (E), or ATOH1 (F) was quantified from sections described in (A–C).

Notably, knocking down both isoforms with sh*-Meis1-all* did not significantly alter the overall proportion of KI67+ or p27+ cells (Figure 4C, D, E and Figure S5D). We interpret this lack of phenotype as resulting from mutual cancellation; the pro-differentiative effect of MEIS1-HdL loss is likely balanced by the anti-differentiative effect of MEIS1-FL loss. However, the specific reduction in the ATOH1-expressing GCP population (Figure 4C, F) may suggest a dominant role for MEIS1-HdL in maintaining this progenitor state, an effect that is not fully compensated for by the simultaneous loss of MEIS1-FL.

### MEIS1-FL and MEIS1-HdL stabilize ATOH1 protein levels

The precise control of ATOH1 protein abundance is critical for regulating the transition of GCPs from proliferation to differentiation. ^8,10,28,29,30^ Our previous work has shown that MEIS1-FL promotes transcription of *Pax6*, activates BMP signaling, and acts to promote ATOH1 degradation. ^16^ However, we found that MEIS1-HdL lacking homeodomain also affects the differentiation of GCPs to GCs (Figure 3, 4), suggesting that MEIS1 proteins may have additional functions to regulate the amount of ATOH1. To test this, we first examined the effect of *Meis1* knockdown on ATOH1 protein levels *in vivo* by measuring ATOH1 fluorescence intensities. In cerebella electroporated with sh-*Meis1-HdL* at P8 and fixed at P11, ATOH1 fluorescence intensities in electroporated cells (GFP+) were lower compared to surrounding non-electroporated cells (Figure S6A, B, E). In contrast, KI67 fluorescence intensities were not significantly affected by sh-*Meis1-HdL* (Figure S6A, B, D). This suggests that MEIS1-HdL may participate in maintaining ATOH1 protein levels in ATOH1-expressing GCPs (Figure S6A, B, D). Similar tendencies were also observed for sh-*Meis1-all* electroporated cerebella (Figure S6A, C–E). However, because sh-*Meis1-all* was designed to suppress both MEIS1-HdL and MEIS1-FL, it was unclear whether MEIS1-FL is also capable of maintaining ATOH1 protein levels at this point.

To further investigate the machinery of MEIS1 isoforms to regulate ATOH1 protein levels, we transfected ATOH1 and MEIS1 expression vectors into N2a cells and performed immunoblotting. We used a GST-tagged ATOH1 (GST-ATOH1) expression vector to discriminate between endogenous and exogenous ATOH1 proteins, although it is known that N2a cells do not express endogenous ATOH1 under normal conditions. ^31^ Administration of MG132, an inhibitor of proteasome-dependent protein degradation, enhanced the ATOH1 signals (Figure S7A, lane 5) relative to the control (Figure S7A, lane 1), consistent with the previous findings that ATOH1 is degraded via proteasome-dependent protein degradation. ^10,11,32,33^ Interestingly, ATOH1 signals were also higher following co-transfection with MEIS1-HdL (lane 3 in Figure S7A) or MEIS1-FL (Figure S7A, lane 4), whereas the N-terminal MEIS1 fragment (1–130 aa) did not exhibit such an effect (Figure S7A, lane 2). To further analyze ATOH1 stability, we immunoblotted transfected N2a cells treated with cycloheximide (CHX) for 0, 4, and 8 h. This revealed that MEIS1-HdL and MEIS1-FL both strongly suppressed ATOH1 degradation to the same extent as MG132 administration (Figure S7 B, C). These observations suggest that both MEIS1-HdL and MEIS1-FL have the capability to suppress the proteasome-dependent degradation of ATOH1.

### MEIS1 inhibits CUL3–mediated ubiquitination of ATOH1

Because the proteasome-dependent degradation is triggered by ubiquitination of proteins, we attempted to assess ATOH1 ubiquitination in cultured cells. N2a cells were co-transfected with GST-ATOH1 and HA-Ub and cultured with MG132 to block proteasome-dependent protein degradation. Then, GST-ATOH1 was pulled down by glutathione beads and immunoblotted with an anti-HA antibody. Consistent with previous findings, ^10,32^ ATOH1 was strongly polyubiquitinated (Figure 5A, lane 1). However, co-transfection with MEIS1-FL or MEIS1-HdL significantly reduced GST-ATOH1 polyubiquitination (Figure 5A, lanes 2 and 3; Figure 5B). Interestingly, MEIS1-HdL suppressed polyubiquitination more strongly than MEIS1-FL (Figure 5B). These findings suggest that both MEIS1-FL and MEIS1-HdL suppress the functions of molecules related to ATOH1 polyubiquitination, eventually blocking ATOH1 degradation. The fact that not only MEIS1-FL but also MEIS1-HdL does this suggests that this function is not elicited by MEIS1-induced transcriptional regulation.

**Figure 5.**
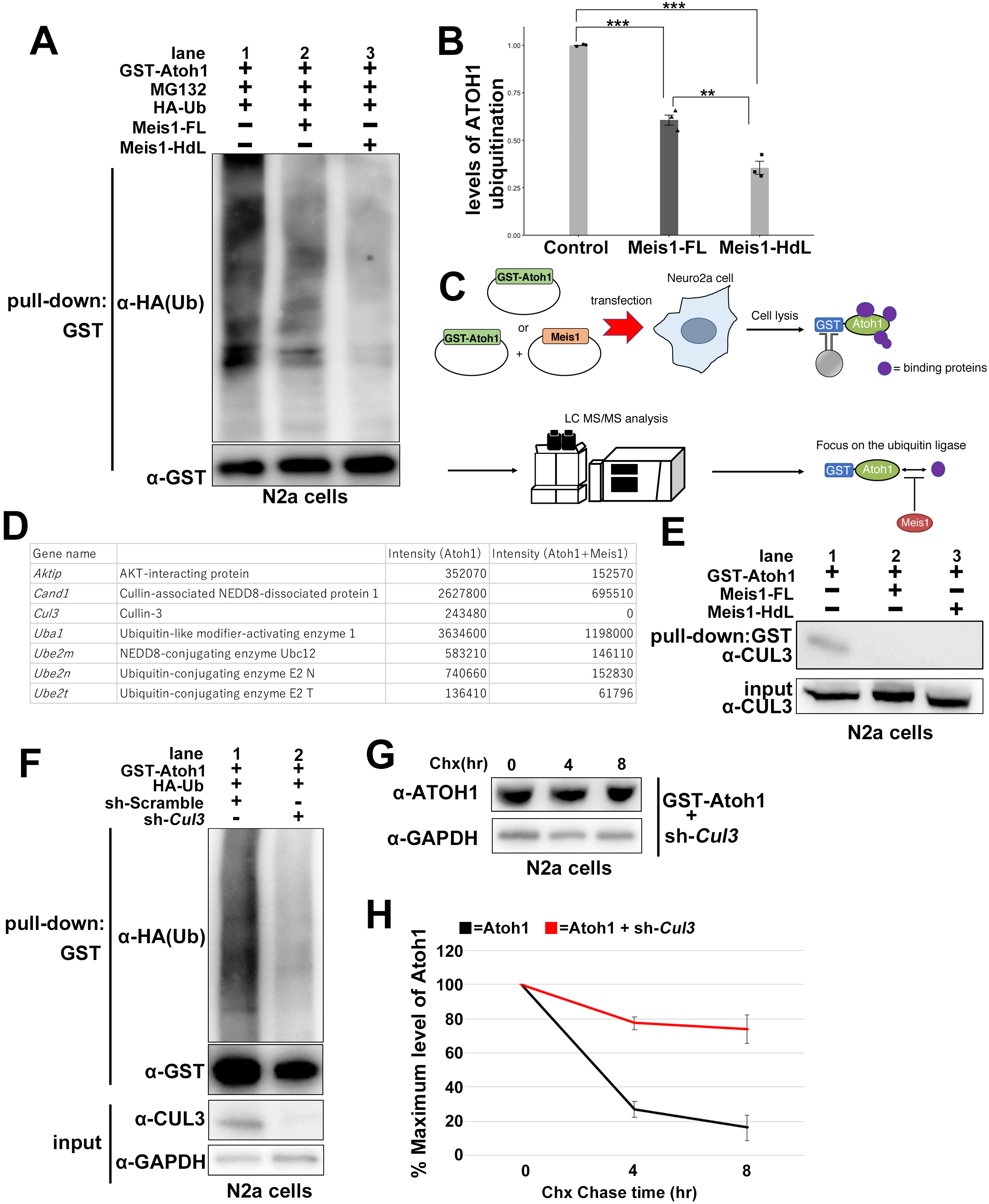
MEIS1 inhibits CUL3-mediated ATOH1 ubiquitination, leading to ATOH1 stabilization. **A.** Immunoblotting of GST pull-down fractions from N2a cells co-transfected with GST-ATOH1, HA-Ub, and either MEIS1-FL or MEIS1-HdL, followed by MG132 treatment for 6 hours. Polyubiquitinated ATOH1 (upper panel) was detected using an anti-HA antibody, while total GST-ATOH1 levels (lower panel) were detected using an anti-GST antibody. **B.** Quantification of polyubiquitinated ATOH1 levels, normalized to total GST-ATOH1 levels from (A). **C.** Schematic illustrating the proteomic analysis strategy to identify ATOH1-binding molecules in the presence or absence of MEIS1 in N2a cells. **D.** List of ubiquitin-related proteins (identified by PANTHER GO analysis) that showed reduced binding to ATOH1 in the presence of MEIS1-FL, as identified by proteomic analysis (as described in C). **E.** Immunoblotting of GST pull-down fractions from N2a cells transfected with GST-ATOH1 alone or co-transfected with GST-ATOH1 and either MEIS1-FL or MEIS1-HdL. Interaction with endogenous CUL3 was determined by immunoblotting with an anti-CUL3 antibody. The interaction is disrupted by the co-expression of either MEIS1-FL or MEIS1-HdL. **F.** Immunoblotting of GST pull-down fractions from N2a cells co-transfected with GST-ATOH1, HA-Ub, and either a control scramble shRNA (sh-Scramble) or a *Cul3* knockdown shRNA (sh-*Cul3*). Polyubiquitinated ATOH1 (upper panel) was detected with an anti-HA antibody, and total GST-ATOH1 levels (middle panel) were detected with an anti-GST antibody. CUL3 expression in input lysates was confirmed by immunoblotting with an anti-CUL3 antibody (lower panel). **G.** Immunoblotting analysis of N2a cell lysates following cycloheximide (CHX) chase assay. N2a cells transfected with GST-ATOH1 and sh-*Cul3* were treated with CHX for 0, 4, and 8 hours. ATOH1 protein levels were detected using an anti-ATOH1 antibody, with GAPDH as a loading control. **H.** Quantification of ATOH1 protein levels from (G), normalized to GAPDH, showing that knockdown of *Cul3* inhibits ATOH1 degradation.

Based on our findings suggesting that MEIS1 isoform proteins affect the function of molecules involved in ATOH1 polyubiquitination, we searched for ATOH1-binding molecules whose binding affinities were altered by the presence of MEIS1 in cultured cells. After transfecting GST-ATOH1 or GST-ATOH1 plus MEIS1-FL into N2a cells, GST-ATOH1 was pulled down, and its binding molecules were subjected to LC–MS/MS (Figure 5C). We obtained many candidate molecules that bind ATOH1 (see Supplementary File for list of binding candidate molecules), identifying several ATOH1-binding molecules whose interaction with ATOH1 was disrupted or much reduced in the presence of MEIS1-FL (Figure 5D). Notably, these included the E3 ubiquitin ligase Cullin-3 (CUL3) and its known partners UBE2M and CAND1 (Figure 5D and Supplementary File for list of binding candidate molecules), ^34,35,36^ suggesting that CUL3 may act as an E3-ligase for ATOH1. To date, HUWE1 is the only E3-ligase reported to participate in ATOH1 degradation. ^10,32^ Although we found that HUWE1 binds to GST-ATOH1, its binding ability was not affected by MEIS1-FL, which further focused our attention on CUL3.

To confirm the proteomic finding that MEIS1 disrupts the ATOH1-CUL3 interaction, we performed co-immunoprecipitation experiments. We co-transfected GST-ATOH1 and MEIS1-FL or MEIS1-HdL into N2a cells, followed by pull-down with GST-ATOH1. Similar to the LC–MS/MS results, immunoblotting showed that CUL3 bound ATOH1 in the absence of MEIS1 (Figure 5E, lane 1), but not in the presence of MEIS1-FL or MEIS1-HdL (Figure 5E, lanes 2, 3).

Next, we examined the involvement of CUL3 in ATOH1 polyubiquitination. GST-ATOH1 and HA-Ub were co-transfected with a control or *Cul3*-knockdown vector (sh-*Cul3*) into N2a cells, which were subjected to pulldown and immunoblotting with the indicated antibodies (Figure 5F), in the absence of MG132. GST-ATOH1 polyubiquitination was strongly suppressed by *Cul3*-knockdown (Figure 5F), suggesting that CUL3 is involved in ATOH1 polyubiquitination. To further analyze the effect of *Cul3 knockdown* on ATOH1 stability, we immunoblotted N2a cells transfected with GST-ATOH1 and control or sh-*Cul3* and administered CHX for 0, 4, and 8 h (Figure 5G). This revealed that *Cul3*-knockdown markedly suppressed ATOH1 degradation in N2a cells (Figure 5H). These observations suggest that CUL3 acts as a key E3 ubiquitin ligase for ATOH1 and that MEIS1-FL and MEIS1-HdL inhibit the interaction between CUL3 and ATOH1, eventually suppressing the polyubiquitination and degradation of ATOH1.

### MEIS1 inhibits the degradation of ATOH1 induced by S328 phosphorylation

Previous studies have shown that phosphorylation of ATOH1 regulates its degradation,^10,32,37^ which led us to the notion that MEIS1 may affect ATOH1 phosphorylation as a part of its stabilization mechanism. To test this, we initially investigated ATOH1 phosphorylation sites by LC–MS/MS using N2a cells transfected with ATOH1 alone or ATOH1 and MEIS1-FL (Figure S8A). We observed that ATOH1 phosphorylation at some sites was increased in the presence of MEIS1-FL (Figure S8B and Supplementary File for list of ATOH1 phosphorylation sites). Particularly, S328 phosphorylation was markedly increased. Consistent with the proteomics data, immunoblot analysis using an antibody against ATOH1 p-S328 indicated that phosphorylation at S328 was much increased in the presence of either MEIS1-FL or MEIS1-HdL (Figure S8C). It has been reported that S328 phosphorylation is involved in the degradation of ATOH1 ^10,32^. Accordingly, the CHX chase assay showed that ATOH1-S328A (Ser 328 of ATOH1 was replaced with Ala; an ATOH1-S328 non-phosphorylated form) was stable, whereas ATOH1-S328D (Ser 328 of ATOH1 was replaced with Asp; an ATOH1-S328 phosphomimic form) was unstable (Figure S8D, F). However, the stability of ATOH1-S328D increased in the presence of MEIS1-FL or MEIS1-HdL (Figure S8E, F). These observations suggest that the MEIS1 isoforms suppress ATOH1 degradation induced by phosphorylation at S328 without inhibiting phosphorylation of S328 ATOH1.

Next, GST-ATOH1 or GST-ATOH1-S328A and HA-Ub were co-transfected into N2a cells; pull-down and immunoblotting were subsequently performed with the indicated antibodies (Figure S8G). The polyubiquitination efficiency of S328A was lower than that of the wild-type ATOH1 (Figure S8G upper panel), which is consistent with the observation that ATOH1-S328A is stable (Figure S8D, F). Furthermore, the interaction of GST-ATOH1-S328A with CUL3 was much weaker than that of GST-ATOH1 with CUL3 (Figure S8G second panel from the top). These findings indicate that S328 phosphorylation is involved in CUL3 binding and polyubiquitination of the ATOH1 protein. MEIS1 isoforms do not suppress S328 phosphorylation but instead act to inhibit the interaction between S328-phosphorylated ATOH1 and CUL3, leading to the stabilization of ATOH1.

### MEIS1 inactivates CUL3 by promoting COP9 signalosome (CSN) and CUL3 complex formation

To understand how MEIS1 inhibits CUL3 activity, we next investigated MEIS1-binding molecules in N2a cells. GST-tagged MEIS1-FL (GST-MEIS1-FL) was transfected into N2a cells, precipitated using glutathione beads, and subjected to LC–MS/MS (Figure S9A). We obtained 1417 MEIS1-binding candidate molecules (see Supplementary File for list of binding candidate molecules). Interestingly, these included several components of the COP9 signalosome (CSN) complex, such as COPS2, COPS3, COPS4, COPS6, COPS7a, and COPS8 (Figure 6A and Supplementary File for list of binding candidate molecules), suggesting that MEIS1-FL interacts with the CSN complex. It is known that the CSN complex, consisting of eight components, suppresses substrate polyubiquitination by the Cullin-family E3 protein ligases. By competitively binding Cullin proteins, the CSN complex masks the substrate-binding regions of Cullin proteins, thus inhibiting their interactions with substrates. ^36^ The CSN complex also suppresses protein neddylation, thereby inhibiting Cullin protein function.

**Figure 6.**
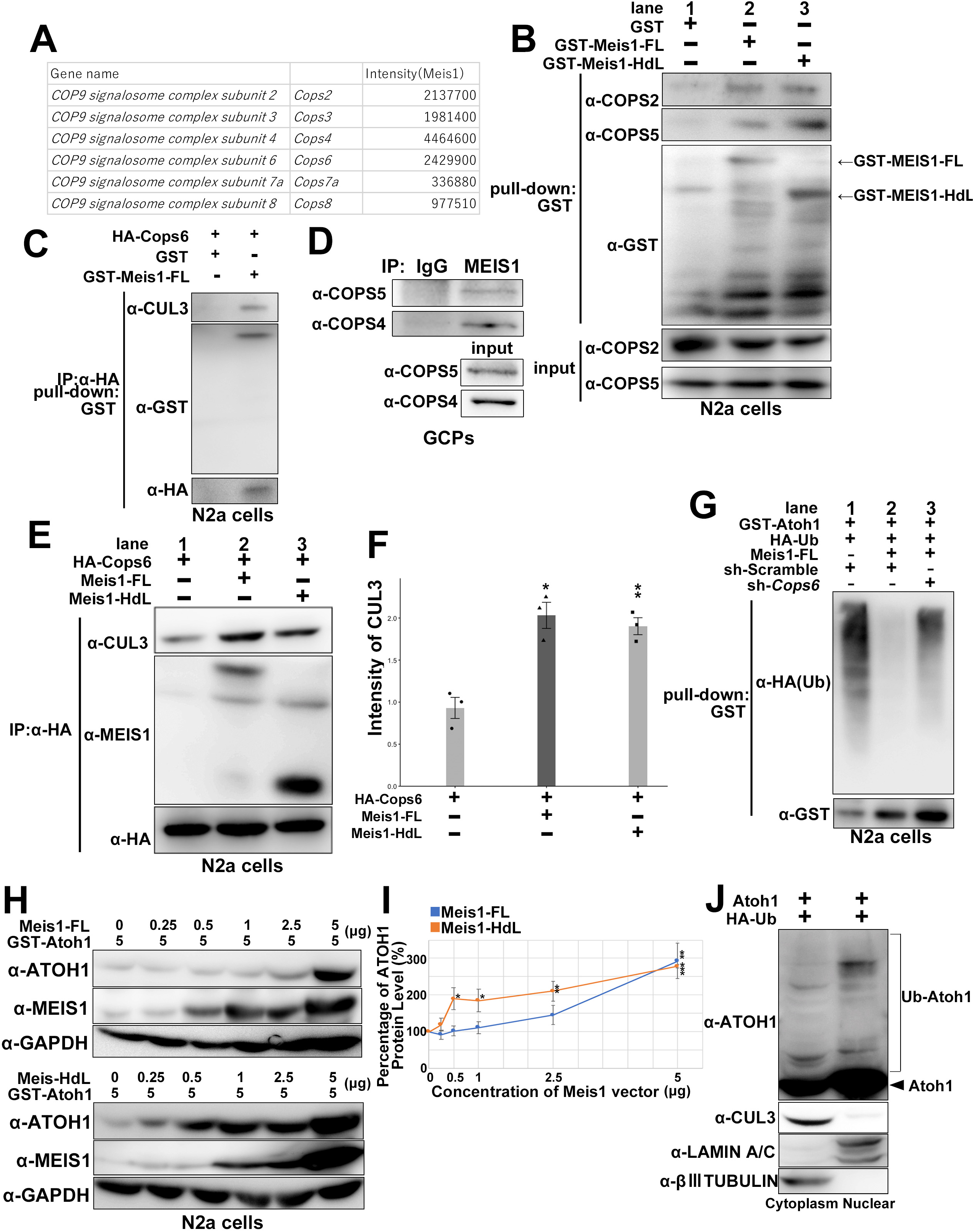
MEIS1 isoforms promote CUL3–COP9 signalosome complex formation to inactivate CUL3. **A.** List of components of the COP9 signalosome (CSN) complex identified as MEIS1-binding partners via proteomic analysis (experimental scheme is shown in Figure S9A). **B.** Immunoblotting of GST pull-down fractions from N2a cells transfected with GST, GST-MEIS1-FL, or GST-MEIS1-HdL. Interactions with endogenously expressed COPS2 and COPS5 were determined using anti-COPS2 and anti-COPS5 antibodies. Total GST-tagged proteins were detected with an anti-GST antibody. **C.** Sequential co-immunoprecipitation assay demonstrating a CUL3–MEIS1–COPS6 tripartite complex. N2a cells were co-transfected with Flag-HA-COPS6 and GST-MEIS1-FL (or GST as control). Lysates were first subjected to immunoprecipitation with an anti-HA antibody, followed by a GST pull-down. The final immunoblot for endogenous CUL3 confirmed its inclusion in a tripartite complex with MEIS1 and COPS6. **D.** Co-immunoprecipitation of endogenous proteins from P7 GCP lysates. Endogenous MEIS1 was immunoprecipitated using an anti-MEIS1 antibody, with IgG serving as a negative control. The resulting precipitates were then immunoblotted for the CSN components COPS4 and COPS5 to detect an interaction. **E.** Co-immunoprecipitation assay to assess the effect of MEIS1 isoforms on the COPS6-CUL3 interaction. Flag-HA-COPS6 was immunoprecipitated using an anti-HA antibody from N2a cells co-expressing a control vector, MEIS1-FL, or MEIS1-HdL. The immunoblots show that co-expression of either MEIS1 isoform enhances the binding of endogenous CUL3 to COPS6. **F.** Quantification of CUL3 signal intensity from (E). **G.** *In vitro* ubiquitination assay demonstrating that COPS6 is required for MEIS1 to inhibit ATOH1 polyubiquitination. N2a cells were co-transfected with GST-ATOH1, HA-Ub, and the indicated constructs (MEIS1-FL, sh-*Cops6*, or controls). Following a GST pull-down, blots were probed for HA (polyubiquitination, top panel) and GST (total ATOH1, bottom panel). The ability of MEIS1-FL to reduce ubiquitination is reversed by the knockdown of *Cops6* (see Figure S9D for knockdown efficiency). **H.** Dose-dependent stabilization of GST-ATOH1 by MEIS1 isoforms. N2a cells were co-transfected with a constant amount of GST-ATOH1 plasmid and increasing amounts (0–5 µg) of plasmids expressing MEIS1-FL or MEIS1-HdL. Cell lysates were immunoblotted for the indicated proteins, with GAPDH as a loading control. **I.** Quantification of ATOH1 protein levels from (H), normalized to GAPDH and plotted against the concentration of the transfected MEIS1 expression vector. The graph illustrates the differential potency of MEIS1-FL and MEIS1-HdL in stabilizing ATOH1. *Asterisks denote a significant difference from the 0 µg control for each respective isoform. **J.** Immunoblot of cytoplasmic and nuclear fractions from N2a cells co-expressing ATOH1 and HA-Ub. The blot shows that while polyubiquitinated ATOH1 is detected in both fractions, CUL3 is predominantly localized to the cytoplasm. βIII-TUBULIN and LAMIN A/C serve as markers for the cytoplasmic and nuclear fractions, respectively.

To confirm the interaction between the CSN complex and MEIS1 isoforms, we transfected GST-MEIS1-FL or GST-MEIS1-HdL into N2a cells, followed by GST pull-down. Immunoblotting revealed that not only MEIS1-FL but also MEIS1-HdL interacted with endogenous CSN components, COPS2 and COPS5 (Figure 6B). We conducted similar pull-down and immunoblotting with HA-tagged COPS6 (HA-COPS6) and GST-MEIS1-truncated mutant (1–130 aa) or GST-MEIS1 isoforms in N2a cells. As expected, HA-COPS6 was found to bind GST-MEIS1-FL and GST-MEIS1-HdL (Figure S9B). In contrast, GST-MEIS1-1–130aa did not bind HA-COPS6 (Figure. S9B). Given that this N-terminal MEIS1 fragment (MEIS1-1–130aa) does not have the ATOH1-stabilizing ability (Figure S7), these observations suggest that binding to the CSN complex is crucial for MEIS1 proteins to stabilize ATOH1.

To test whether MEIS1, CSN complex and CUL3 form a tripartite complex, we performed a sequential immunoprecipitation experiment. HA-COPS6 and GST-MEIS1-FL were transfected into N2a cells, followed by immunoprecipitation with HA and successive pull-down with GST. Immunoblotting revealed a significant signal for endogenous CUL3 (Figure 6C), suggesting that the CSN complex, MEIS1-FL, and CUL3 form a tripartite complex in cells. To test the interaction of endogenously expressed MEIS1 and CSN complex, immunoprecipitation was performed on the GCP lysates with the MEIS1 antibody recognizing both MEIS1 isoforms. Immunoblotting with COPS4 or COPS5 suggested that endogenous MEIS1 and the CSN complex interact with each other (Figure 6D). Given that MEIS1-HdL interacts with components of the CSN complex in N2a cells (Figure 6B, Figure S9B), it was suggested that not only MEIS1-FL but also MEIS1-HdL interacts with the CSN complex in GCPs.

We next tested whether the presence of MEIS1 isoforms affects the CUL3-CSN interaction. N2a cells were transfected with HA-COPS6 and MEIS1-FL or MEIS1-HdL, followed by immunoprecipitation with the HA antibody. Immunoblotting showed that the amount of co-precipitated CUL3 was significantly greater in the presence of either MEIS1-FL or MEIS1-HdL, compared to control (Figure 6E, F). This suggests that MEIS1 isoforms inactivate CUL3 by enhancing its association with the CUL3-inhibitory complex, CSN. We further tested the possibility that CUL3 is required for the binding between MEIS1 isoforms and the CSN complex. However, it was not likely because the introduction of sh-*Cul3* into N2a cells did not affect the binding between HA-COPS6 and MEIS1-FL or MEIS1-HdL (Figure S9C). To investigate whether the CSN complex is involved in the function of MEIS1 to suppress ATOH1 polyubiquitination, GST-ATOH1 and HA-Ub were transfected into N2a cells in the presence of MG132, followed by GST pull-down. Immunoblotting with HA revealed that MEIS1-FL introduction reduced ATOH1 polyubiquitination (Figure 6G, lanes 1 and 2). However, co-introduction of the knockdown vector for *Cops6* (sh-*Cops6*, Figure S9D) ameliorated the effects of MEIS1-FL (Figure 6G, lane 3). These findings suggest that MEIS1 suppresses ATOH1 polyubiquitination and degradation by promoting the CSN complex–CUL3 interaction and by inhibiting the CUL3–ATOH1 interaction. When transfected into N2a cells, the expression of NRF2, NCOA3, and PP2A/C was much higher in the presence of MG132 (Figure S9E, lane 4) than in the control (Figure S9E, lane 1), consistent with previous findings that these proteins are degraded by CUL3-dependent polyubiquitination. ^38,39,40^ Co-introduction of MEIS1-FL or MEIS1-HdL strongly increased NRF2 and PP2A/C expression and moderately increased that of NCOA3 (Figure S9E, lanes 2 and 3), relative to the control (Figure S9E, lane 1). This suggests that these MEIS1 isoforms may broadly suppress CUL3-dependent degradation of various proteins.

Transfected tagged COPS6 was expressed in both intracellular regions in N2a cells (Figure S9F), consistent with a previous report that the CSN complex localizes in both the cell nucleus and cytoplasm. Similar COPS6 subcellular localization was observed following co-transfection with MEIS1-FL or MEIS1-HdL (Figure S9G, H), suggesting that neither MEIS1-FL nor MEIS1-HdL affects CSN-complex subcellular localization.

Thus, MEIS1-FL and MEIS1-HdL have similar functions in that they both inhibit the degradation of ATOH1. However, there may be differences in the strength of their effects. To test this, GST-ATOH1 was co-transfected with MEIS1-FL or MEIS1-HdL, using the indicated quantities of DNA (0–5 μg, Figure 6H). While large amounts of MEIS1-FL vector were required to suppress ATOH1 degradation, this was achieved using very small amounts of MEIS1-HdL vector (Figure 6H, I). This suggests that MEIS1-HdL suppresses ATOH1 degradation more effectively than MEIS1-FL. Fractionation using N2a cells transfected with ATOH1 and HA-Ub revealed that polyubiquitinated ATOH1 was localized in both the cytoplasm and nucleus (Figure 6J). In contrast, CUL3 was localized abundantly in the cytoplasm and negligibly in the nuclei (Figure 6J), consistent with prior suggestions. ^41^

These findings suggest that ATOH1 is polyubiquitinated by CUL3 in the cytoplasm, probably just after ATOH1 protein synthesis, while ATOH1 polyubiquitination in nuclei is elicited by other E3 ligases. MEIS1-HdL was localized exclusively in the cytoplasm, whereas MEIS1-FL was localized predominantly in the nucleus (Figure S2B, C). This difference in intracellular localization between MEIS1-FL and MEIS1-HdL probably accounts for the differential efficacy in suppressing CUL3-dependent ATOH1 degradation.

### The MEIS1-HdL–CSN pathway maintains GCPs in an immature state

We showed that MEIS1-HdL introduction into the EGL suppressed the differentiation from GCPs to GCs: 3 d after electroporation, the proportions of Ki67^+^ and ATOH1^+^ GCPs were higher, while those of p27^+^ GCs were lower (Figure 3C–F). On the other hand, our *in vitro* experiments revealed that ATOH1 stabilization by MEIS1 was mediated by the CSN complex (Figure 6G). To link these *in vivo* and *in vitro* findings, we next tested whether the CSN complex is required for the *in vivo* function of MEIS1-HdL. We co-electroporated sh-*Cops6* together with the MEIS1-HdL into the P8 EGL under the experimental conditions shown in Figure 3C. The effects of MEIS1-HdL were cancelled by co-introduction with sh-*Cops6* to the same extent as sh-*Meis1-HdL* single introduction (Figure 7A–E). This suggests that CSN is required for MEIS1-HdL to function in GCPs, and that the CSN complex works with MEIS1-HdL to suppress the GCP differentiation into GCs, during cerebellar development.

**Figure 7.**
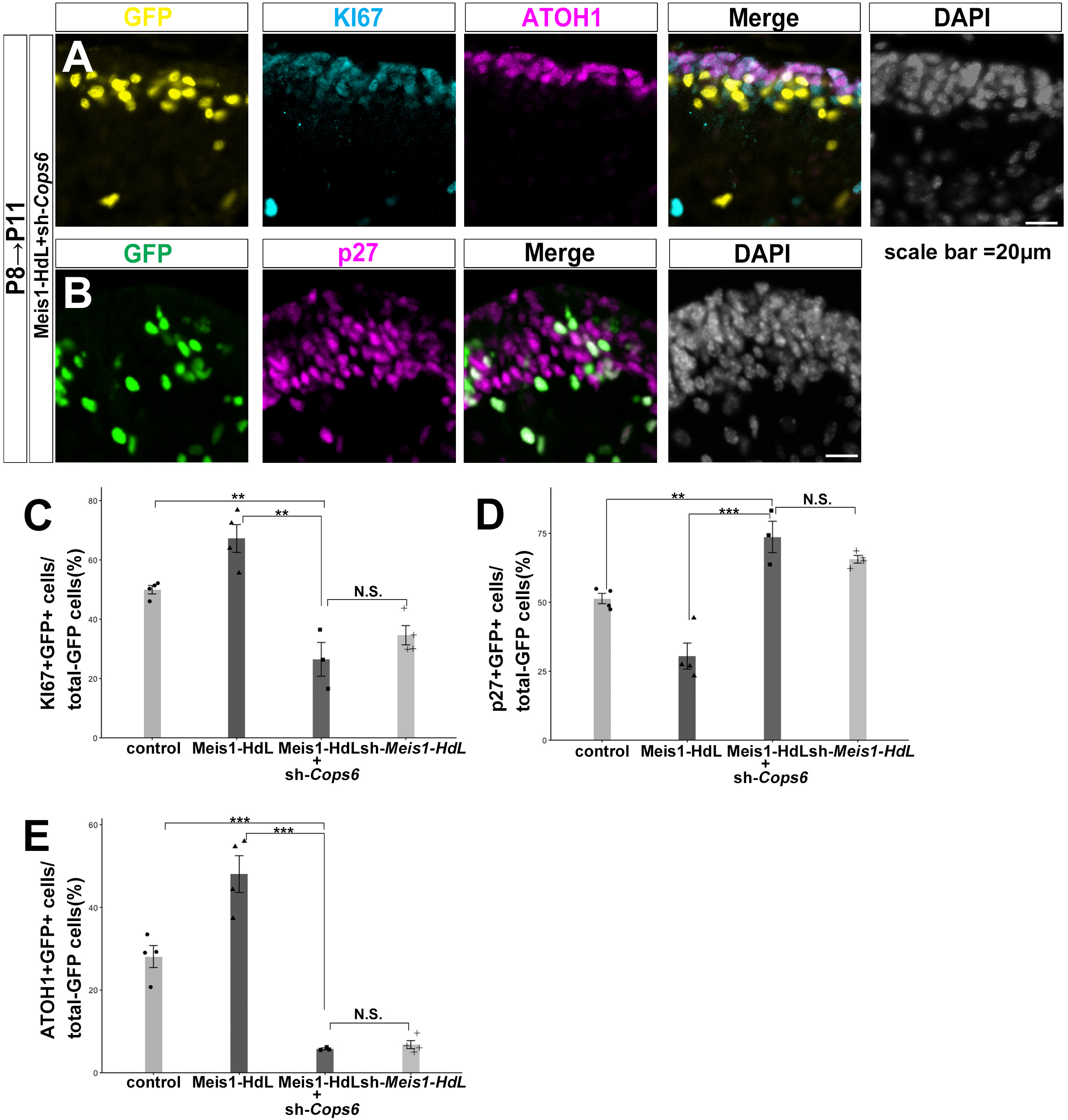
The COP9 signalosome is required for MEIS1-HdL to maintain GCPs in an undifferentiated state. **A.** Representative immunofluorescence images of P11 cerebella following *in vivo* electroporation at P8. Sections were immunostained for ATOH1 (magenta) and KI67 (cyan). Electroporated cells are identified by co-electroporated H3.1-EGFP (yellow). These images show GCPs electroporated with MEIS1-HdL and sh-*Cops6*. **B.** Representative immunofluorescence images of P11 cerebella following *in vivo* electroporation at P8. Sections were immunostained for p27 (magenta). Electroporated cells are identified by co-electroporated H3.1-EGFP (green). These images show GCPs electroporated with MEIS1-HdL and sh-*Cops6*. **C–E.** Quantification of electroporated cell differentiation status. The percentage of GFP+ cells expressing KI67 (C), p27 (D), or ATOH1 (E) was quantified from cerebellar sections.

## Discussion

The dynamic switching between isoforms through alternative splicing and alternative transcriptional start site is increasingly recognized as a sophisticated strategy for achieving complex cellular control during neural development. ^4,5^ While recent long-read RNA-seq technology has revealed the complexity of isoforms, the molecular mechanisms by which these diverse isoforms control complex neurodevelopment remain largely unknown. In this study, we used GCPs as model cells and *Meis1* as a model gene to clarify how the proliferation and differentiation of neural progenitor cells are precisely controlled by multiple isoforms produced from a single gene.

Our long-read RNA-seq and immunoblotting revealed that the *Meis1* gene produces two protein isoforms in GCPs: MEIS1-FL, which has a homeodomain, and MEIS1-HdL, which lacks the homeodomain. *In vivo* electroporation revealed that these two isoforms have opposite effects on GCP proliferation and differentiation. MEIS1-FL inhibits the proliferation of GCPs and promotes differentiation, whereas MEIS1-HdL promotes their proliferation and inhibits differentiation. We have previously reported the function of the former, MEIS1-FL. ^16^ Importantly, the conditional knockout allele used in that study targets an exon specific to homeodomain-containing isoforms, leaving the MEIS1-HdL isoform intact. MEIS1-FL localizes to the nucleus of GCPs and GCs and promotes *Pax6* transcription. Then, the transcription factor PAX6 promotes differentiation from GCPs to GCs by activating *Smad1* transcription and promoting BMP signal-dependent degradation of ATOH1.

Our cell culture experiments revealed that both MEIS1-HdL and MEIS1-FL have functions other than transcription factor activity. These two isoforms have the ability to suppress CUL3-dependent polyubiquitination and degradation of ATOH1. However, it was shown that MEIS1-HdL has a much stronger ATOH1-stabilization ability than MEIS1-FL. This can be explained by differences in subcellular localization. Since CUL3 is mainly localized in the cytoplasm, CUL3-dependent ATOH1 degradation occurs in the cytoplasm. While MEIS1-FL is localized in the nucleus, MEIS1-HdL resides in the cytoplasm. Therefore, it is thought that MEIS1-HdL can more efficiently inhibit CUL3-dependent ATOH1 degradation than MEIS1-FL. In line with this notion, although MEIS1-FL shows partial cytoplasmic localization when overexpressed *in vitro*, it is largely absent from the cytoplasmic fraction of GCPs, indicating that MEIS1-HdL serves as the major cytoplasmic isoform physiologically. Furthermore, MEIS1-HdL is expressed only in undifferentiated, proliferative GCPs and not in GCs. From these findings, it is suggested that MEIS1-HdL promotes the proliferation of GCPs and inhibits their differentiation by suppressing the CUL3-dependent ATOH1 degradation in the cytoplasm, and that the loss of Meis1-HdL expression in GCPs may be an important trigger for their differentiation into GCs.

Thus, MEIS1-FL and MEIS1-HdL have opposite effects on ATOH1 degradation and GCP differentiation, and their expression timing is also contrasting. During cerebellar development, MEIS1-FL is continuously expressed from GCPs to GCs, whereas MEIS1-HdL is only expressed in immature GCPs. In immature GCPs where both are co-expressed, the action of MEIS1-HdL is dominant, leading to the stabilization of the ATOH1 protein. However, as development progresses, MEIS1-FL becomes predominant due to the loss of MEIS1-HdL expression, resulting in the degradation of ATOH1 and progression of differentiation into GCs. Thus, these two MEIS1 isoforms are good examples of a strict control mechanism of neurogenesis by antagonistic isoforms from a single gene.

We identified CUL3 as a novel E3 ligase for ATOH1 and discovered that MEIS1 suppresses its function. It is known that CUL3 activity is suppressed by a large complex called the COP9 signaling complex (CSN). ^42^ Previously, it has been shown that the low-molecular-weight compound, inositol hexakisphosphate (IP6), and its phosphorylating enzyme, protein IP6K1, regulate the binding of CULLIN family proteins and CSN. ^43,44^ This study suggests that MEIS1, especially the cytoplasm-localized MEIS1 isoform (MEIS1-HdL), has a similar function to enhance the binding between CUL3 and CSN, thereby suppressing CUL3 activity, eventually leading to elaborate control of ATOH1 degradation. Furthermore, we showed that MEIS1 has the ability to stabilize CUL3 target proteins other than ATOH1, such as NRF2 and PP2A/C. Moreover, our proteome analyses suggested that MEIS1 binds to CULLIN family proteins other than CUL3, such as CUL1 and CUL4. This raises the possibility that MEIS1 is involved in the degradation control of a wide range of proteins.

These findings have broad biological significance beyond the context of cerebellar development. The dysregulation of key developmental transcription factors like MEIS1 as well as the Cullin-RING E3 ubiquitin ligase machinery, including CUL3 and CSN, is often found in various tumors. ^45^ For example, MEIS1 and COPS5 (a component of CSN) are overexpressed in some acute myeloid leukemia. ^46,47^ Transgenic mice overexpressing COPS5 show an enlarged pool of hematopoietic stem cells, exhibiting a myeloproliferative disorder-like phenotype. ^48^ Similarly, overexpression of MEIS1 enhances the proliferation of hematopoietic stem cells. ^49^ Until now, MEIS1 has been thought to cause leukemia and several types of tumors through its transcription regulatory function. However, this study suggests that MEIS1 may also contribute to tumorigenesis by controlling the degradation of relevant proteins through its non-transcriptional regulatory function.

A well-known example of precise functional control by multiple isoforms is the NMDA-type glutamate receptor, known to be involved in learning and memory. The *Glun1* gene, which encodes the essential subunit, GLUN1, of this receptor, produces several isoforms through alternative splicing. ^50^ GLUN1 isoforms with different N-terminal sequences cause differences in the opening and closing speeds of NMDA receptor channels. ^51^ In addition, GLUN1 isoforms with distinct C-terminal sequences result in differences in the intracellular localization of NMDA receptors. ^52^ These isoforms are strictly regulated for expression timing during development and are essential for normal neural circuit formation. ^53^ This example also suggests that multiple isoforms derived from individual genes strictly and precisely control the development and function of the nervous system with their distinct molecular functions, intracellular localization, and expression profiles. Further research combining long-read sequencing and single-cell analysis technologies will reveal a higher-resolution blueprint of the brain, showing which isoforms are expressed by individual cells and at what timing.

## Materials and Methods

### Animals

All animal experiments in this study were approved by the Animal Care and Use Committee of the National Institute of Neuroscience, National Center of Neurology and Psychiatry (Tokyo, Japan; project 2019028R2). Mice were housed in SPF with food and water ad libitum. ICR pups were obtained from SLC (Japan). *Atoh1^AtGFP/+^* (The Jackson Laboratory: B6.129S-Atoh1tm4.1Hzo/J /J, Stock No: 013593) and *Ki67^KiRFP/+^* (The Jackson Laboratory: Mki67tm1.1Cle/J /J, Stock No: 029802) mice were obtained from The Jackson Laboratory.

### Long-Read cDNA Sequencing Sample Preparation

GCP isolation was performed as previously described. ^54, 55^ For the long-read sequencing samples, isolated P7 GCPs were plated and cultured for 48 h in the presence of 200 nM smoothened agonist. Following the culture period, total RNA was extracted using the Direct-zol™ RNA MiniPrep Plus kit (Zymo Research). For Nanopore sequencing, cDNA libraries were prepared from 50 ng of total RNA using the SMART-Seq® v4 Ultra® Low Input RNA Kit for Sequencing (Takara Bio USA, Inc.) according to the manufacturer’s protocol, with 16 PCR cycles for cDNA amplification. Custom primers were utilized for cDNA synthesis:

3’ SMART-Seq CDS Primer II A: AAGCAGTGGTATCAACGCAGAGTACTTTTTTTTTTTTTTTTTTTTTTTTTTTTTT VN (HPLC-purified)

PCR Primer II A: AAGCAGTGGTATCAACGCAGAGT (HPLC-purified)

The synthesized cDNA was then sequenced on a Nanopore PromethION platform.

### Long-Read cDNA Sequencing Data Analysis

Raw FASTQ files obtained from Nanopore PromethION sequencing were processed using the IsoQuant^56^ pipeline for alignment, transcript quantification, and novel isoform identification. Alignment was performed within the IsoQuant pipeline using Minimap. ^57^ The quality of the resulting sequencing data was assessed prior to analysis. The reads exhibited a high average quality score of approximately 30 (Figure S1A), and the read length distribution peaked around 1000 base pairs (Figure S1B), confirming the high quality and robust coverage of the dataset. The resulting GTF files, containing quantified and identified transcripts, were then subjected to curation using Sqanti3. ^58^

### RT-PCR

RNA was extracted from isolated GCP using Direct-zolTM RNA MiniPrep w/TriReagent (ZYMO RESEARCH). The ReverTra Ace® qPCR RT Kit (TOYOBO) was used to generate cDNA. The primer sequences used are as following:

Meis1-FL: CACAAAAAGCGTGGCATCTT and TGATGCCCATGTGCTGCTGA, Meis1-HdL: ATGGCGCAAAGGTACGACGA and TCAGAAGGGTAAGGGTGCTTGC.

### Quantitative PCR (qPCR)

Isolated GCPs were suspended in cold DPBS containing 1% BSA. Cell sorting was performed in FACSAriaTM Fusion. Gating of GFP and RFP negative populations was performed using GCPs from WT mice. Fluorescence compensation of RFP and GFP was performed using GCP of *Ki67^KiRFP/+^*or *Atoh1^AtGFP/+^* mice. Cells were sorted into cold DPBS containing 1% BSA and centrifuged at 800 × g for 3 min.

cDNA was generated with ReverTra Ace® qPCR RT Kit (TOYOBO). Relative gene expression was compared with the geometric mean of 18s rRNA. The primer sequences used are as following:

18s rRNA: CCCGAAGCGTTTACTTTGAA and CCCTCTTAATCATGGCCTCA, Meis1-FL: ACAGCAGTGAGCAAGGTG and CAGAAGGGTAAGGGTGTGTT, Meis1-HdL: TGAGCAAGCACCCTTACCCTTCTGAA and GACTGCTCGGTTGGACTG, Atoh1: AGCTTCCTCTGGGGGTTACT and TTCTGTGCCATCATCGCTGT.

### Plasmids

The expression vectors of Atoh1, Meis1 1-130aa and Meis1-FL were cloned from cDNAs from C57/BL6 mouse cerebellum. Cloned fragments were inserted into a pCAGGS vector or pEF-BOS-GST vectors (gifts from K. Kaibuchi, Fujita Health University, Nagoya, Japan) vector and cloned sequences confirmed by sequencing. pCAG-H3.1-EGFP vector was a gift from Dr. N. Masuyama. HA-Ub vector was kindly provided by Dr. S. Wakatsuki (NCNP, Tokyo, Japan). Flag-HA-Cops6 was purchased from addgene (Plasmid #22542). Meis1-HdL, Res-Meis1-HdL and mutated forms of Atoh1 S328A and S328D were generated following the mutagenesis protocol of PrimeStarMax (Takara). The primers we used to create Res-Meis1-HdL were GAGCAGGCCCCTTTACCCTTCTGAAGAACA and TAAAGGGGCCTGCTCACTGCTGTTATCCCC. shRNAs were generated by inserting the double-stranded oligonucleotides into a mU6 pro vector. The targeting sequence of each shRNAs was designed by siDirect 2.0(reference) and the target sequences are indicated below.

sh*-Meis1-HdL*: 5′- CAGTGAGCAAGCACCCTTA-3′,

sh-*Meis1-all*: 5′- GCACAAGATACAGGACTTACC -3′ (reference),

sh-*Cul3*: 5′- AGCTGCTATAGTGCGAATAAT -3′,

sh-*Cops6-1*: 5′-ACCAAGGAGGAGCAGTTTAAA -3′,

sh-*Cops6-2*: 5′-TTGAGTCTGTCATCGATATAA -3′.

### Cell culture, transfection, and drug treatment

N2a cells were obtained from ATCC (Manassas, VA, USA) and cultured in Dulbecco’s modified Eagle medium (DMEM) containing 10% fetal bovine serum (FBS) and 100 U/ml penicillin–streptomycin. Transfection of N2a cells was performed using transfection reagent (Bio-Rad, Hercules, CA, USA).

Cycloheximide (100 μM) was diluted in dimethyl sulfoxide (DMSO) and added to N2a cells, as indicated. MG132 diluted in DMSO was administered to the cultured cells indicated, after which cells were harvested at the indicated times.

### Immunohistochemistry

Tissues were fixed with 4% paraformaldehyde (PFA) in PBS and cryoprotected with 30% sucrose in PBS. After tissues were embedded in optimal cutting temperature (OCT) compound, cryosections were made at 16 μm. Sections were incubated in blocking buffer containing 1% BSA and 0.1% Triton X-100 in PBS at room temperature (RT between 20 and 25°C) for 1 h and subsequently immunolabeled using the following primary antibodies in blocking buffer at 4 °C overnight. Specimens were subsequently rinsed with PBS and incubated with secondary antibodies conjugated with Alexa Fluor 488, Alexa Fluor 568, Alexa Fluor 594, or Alexa Fluor 647 (1:400; Invitrogen) and DAPI (1:3,000; Invitrogen) in blocking buffer containing 1% BSA and 0.2% Triton X-100 in PBS at RT for 2 h. Fluorescence images were acquired using a Zeiss LSM 780 confocal microscope system (Carl Zeiss) and ZEN 2009 software. Quantification of the fluorescence intensity of immunolabeled cells was performed using the “Measure” functions of ImageJ. Antibodies used in this study were as follows: anti-ATOH1 antibody (homemade, rabbit), anti-KI67 antibody (eBioscience, 14-5698-82, rat), anti-p27 antibody (MBL, 554, rabbit), anti-HA(C29F4) (CST, 3724S, rabbit), anti-GFP antibody (kind gift from Dr. A. Imura, Foundation for Biomedical Research and Innovation, Kobe, Japan rat), anti-GFP (Aves Labs, GFP-1010, chicken).

### *In vivo* electroporation

*In vivo* electroporation in neonatal mice has been described previously. ^16^ Expression plasmids were diluted to 1 μg/μl, shRNAs to 2 μg/μl, and pCAG-H2BGFP to 0.5 μg/μl in Milli-Q water (Millipore). Fast Green was added to visualize the plasmid solutions, which were injected into P8 ICR cerebella over the skull. Mice received electric pulses (80 V for 50 ms, with 150 ms intervals) using forceps-type electrodes (NEPA Gene, Chiba, Japan). The pups were kept warm at 37 °C during recuperation and returned to the litter after fully recovering. Pups were fixed with 4% PFA for 3 days after electroporation.

### Immunoblotting

Proteins were transferred to polyvinylidene fluoride membranes, which were incubated with primary antibodies at 4 °C overnight. After the membranes were incubated with secondary antibodies at room temperature for 2 h, horseradish peroxidase substrate (Millipore) was applied, and immuno-signals were detected using a LAS4000 system (Fujifilm) and FUSION SOLO S (Vilber). Antibodies used in this study were as follows: anti-GAPDH (CST, #2118, rabbit), anti-GST (CST, #2622, rabbit), anti-HA (CST, #3724, rabbit), anti-ATOH1 (homemade, rabbit), anti-MEIS1-HdL antibody (homemade, rabbit), anti-MEIS1 (abcam, ab19867, rabbit), anti-MEIS1(MyBioSource, MBS9403588, rabbit), anti-CUL3 (CST, #2759, rabbit), anti-COPS2 (Bethyl Laboratories, A300-028A, rabbit), anti-COPS5 (Bethyl Laboratories, A300-014A, rabbit), anti-SRC-3 (NCOA3; CST, #2126, rabbit), anti-PP2A/C subunit (CST, #2038, rabbit), anti-NRF2 (Novusbio, NBP1-32822, rabbit), anti-LAMIN A/C (CST, #4777, mouse), anti-βIIITUBULIN (Millipore, MAB1637, mouse), and anti-ATOH1 p-S328 (homemade, rabbit).

### GST pull-down assay, protein purification, and phospho-MS

GST-tagged vectors expressing Atoh1, Atoh1 (S328A), Atoh1(S328D), Meis1-FL, and Meis1-HdL were transfected into N2a cells, which were cultured for 2 days. Cells were harvested, as described, and after lysis, supernatants were incubated with glutathione-conjugated Sepharose beads (GE Healthcare) at 4 °C overnight. Sepharose beads were washed twice and resuspended in sample buffer. For protein purification, washed sepharose beads were incubated with TED buffer containing 150 mg of reduced-form glutathione (pH 7.4) at 4 °C for 30 min, and the supernatants were stored at -80 °C. Phospho-MS was performed, as previously described. ^59^

### Statistical analyses

Pairwise comparisons between the means of different groups were performed using a Student’s t-test (two-tailed, unpaired). The difference between two subsets of data was considered statistically significant if the Student’s t test gave a significance level of *p < 0.05, **p < 0.01, or ***p < 0.001. The data are reported as the mean ± SEM.

## Supporting information

Supplementary File for the high diversity isoform list.

Supplementary File for list of ATOH1 phosphorylation sites

Supplementary File for list of binding candidate molecules

## Data and Code Availability

The long-read cDNA sequencing data generated in this study have been deposited in the DNA Data Bank of Japan (DDBJ) under accession number PRJDB15106. The accession numbers for the individual BioSamples are SAMD00572392, SAMD00572393, and SAMD00572394.

The full list of proteins identified by proteomic analysis and the list of genes with high isoform diversity are provided as a Supplementary File for the high diversity isoform list.

This paper does not report original code.

## Acknowledgments

We thank all members of Department of Biochemistry and Cellular Biology for fruitful comments and discussion.

This work is supported by Japan Society for the Promotion of Science KAKENHI (22H04925(PAGS), 22K15211 to T.O. and 22H02730, 25K02372 to M.H.), AMED (JP24wm0425005h0004, JP25ek0109764h0002 to M.H.), an Intramural Research Grant of NCNP (4-6, 6-9, 7-8 to M.H., 7-9 to S.M.), Japan Health Research Promotion Bureau (2020-B-07, 2024-D-01 to M.H.), Multilayered Stress Diseases (JPMXP1323015483) Science Tokyo to M.H., and Tokumori Yasumoto Memorial Trust to M.H.

## Author contributions

Conceptualization: T.O. and M.H. Methodology: T.O. Investigation: T.O., T.A., R.S., K.I., K.J., M.M., K.S., K.N., I.H., D.K., T.N., S.T., Y.S., K.K., S.M., and M.H. Formal Analysis: T.O. Resources: D.K. Writing—original draft: T.O. and M.H. Writing – review and editing: all authors Visualization: T.O. Funding acquisition: T.O. and M.H. Supervision: T.O., S.M. and M.H.

## Supplemental Figure Legends

**Figure S1.**
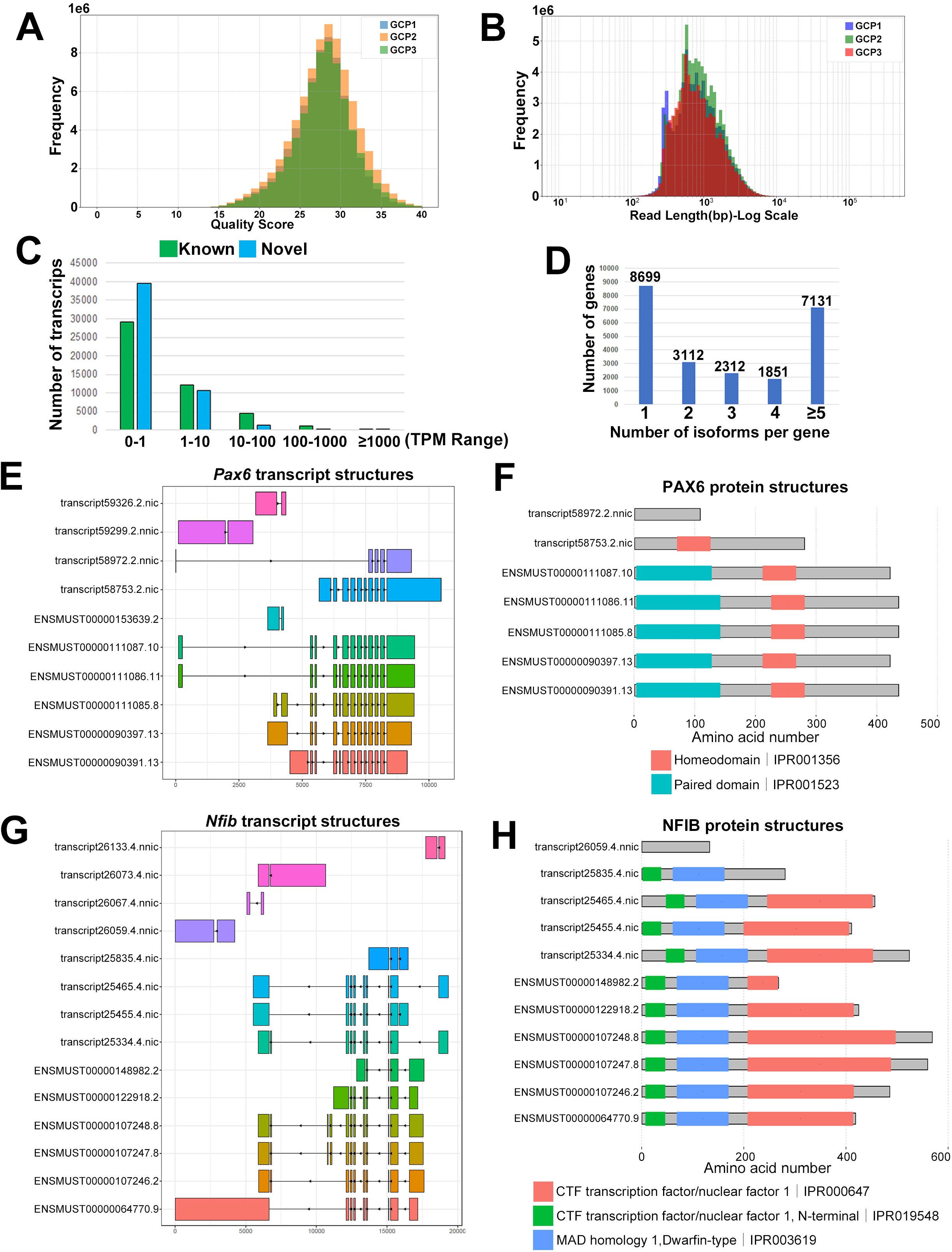
Quality control and overview of long-read cDNA sequencing data. **A.** Histogram showing the distribution of read quality scores (Q-scores) for Nanopore long-read cDNA sequencing data from GCP samples. **B.** Histogram showing the distribution of read lengths for Nanopore long-read cDNA sequencing data from GCP samples. **C.** Histograms showing the distribution of expression levels (Transcripts Per Million, TPM) for known transcripts (left) and novel transcripts (right). **D.** Distribution of the number of isoforms per gene, including all identified transcripts without TPM filtering. The graph shows the percentage of genes binned by their isoform count (1, 2, 3, 4, or ≥5). **E.** Transcript isoform structures of *Pax6*, generated using ggtranscript. **F.** Predicted protein domain structures of coding PAX6 isoforms. Protein sequences were obtained via Sqanti3, with domains predicted by InterPro and visualized using drawProteins. Key conserved domains are shown. Some isoforms lack the paired domain but retain the homeodomain, and a short isoform lacks all canonical domains. **G.** Transcript isoform structures of *Nfib*, generated using ggtranscript. **H.** Predicted protein domain structures of coding *Nfib* isoforms. Protein sequences were obtained via Sqanti3, with domains predicted by InterPro and visualized using drawProteins. Some isoforms contain large deletions, including a short variant that lacks all predicted domains.

**Figure S2.**
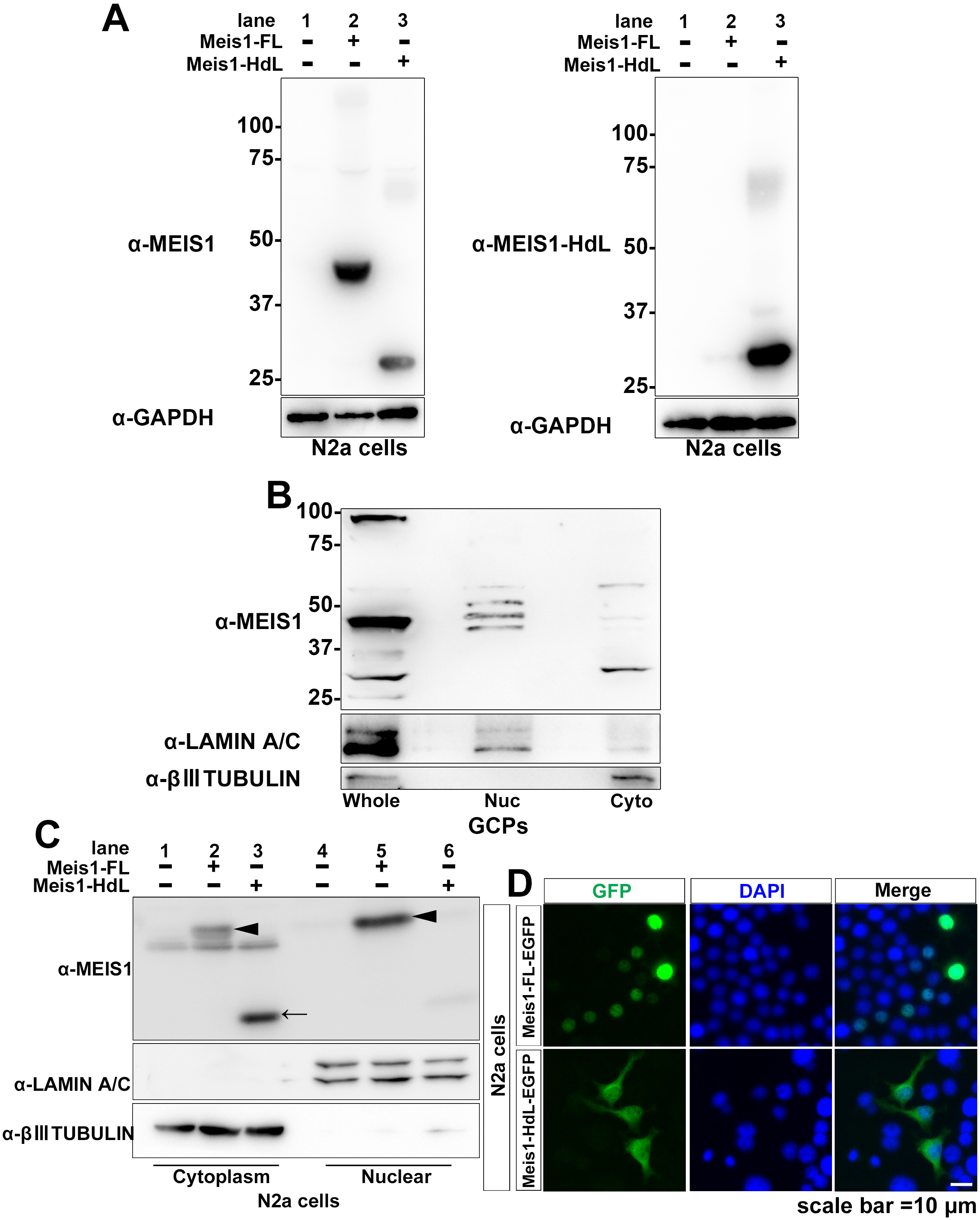
Validation of MEIS1-HdL-specific antibody and further confirmation of MEIS1 isoform subcellular localization. **A.** Validation of the MEIS1-HdL-specific antibody by immunoblot. Lysates from untransfected N2a cells or cells expressing MEIS1-FL or MEIS1-HdL were analyzed. The left panel, probed with a pan-MEIS1 antibody, detects both isoforms. The right panel shows that the newly generated antibody specifically recognizes MEIS1-HdL and not MEIS1-FL. **B.** Subcellular localization of endogenous MEIS1 isoforms in P7 GCPs, analyzed by immunoblotting of subcellular fractions. The blot shows that MEIS1-FL is predominantly nuclear, while MEIS1-HdL is primarily cytoplasmic. LAMIN A/C and βIII-TUBULIN were used as nuclear and cytoplasmic fraction markers, respectively. **C.** Immunoblot analysis showing the subcellular localization of overexpressed MEIS1 isoforms in N2a cells. Following transfection with untagged MEIS1-FL or MEIS1-HdL, cytoplasmic and nuclear fractions were blotted for MEIS1. The results confirm the predominantly nuclear localization of MEIS1-FL (arrowheads) and cytoplasmic localization of MEIS1-HdL (arrows). LAMIN A/C and βIII-TUBULIN serve as nuclear and cytoplasmic fraction markers, respectively. **D.** Representative immunofluorescence images showing subcellular localization of MEIS1-FL-EGFP and MEIS1-HdL-EGFP in N2a cells. Cells were transfected with MEIS1-FL-EGFP or MEIS1-HdL-EGFP expression vectors and stained for GFP (green) and DAPI (blue, for nuclei).

**Figure S3.**
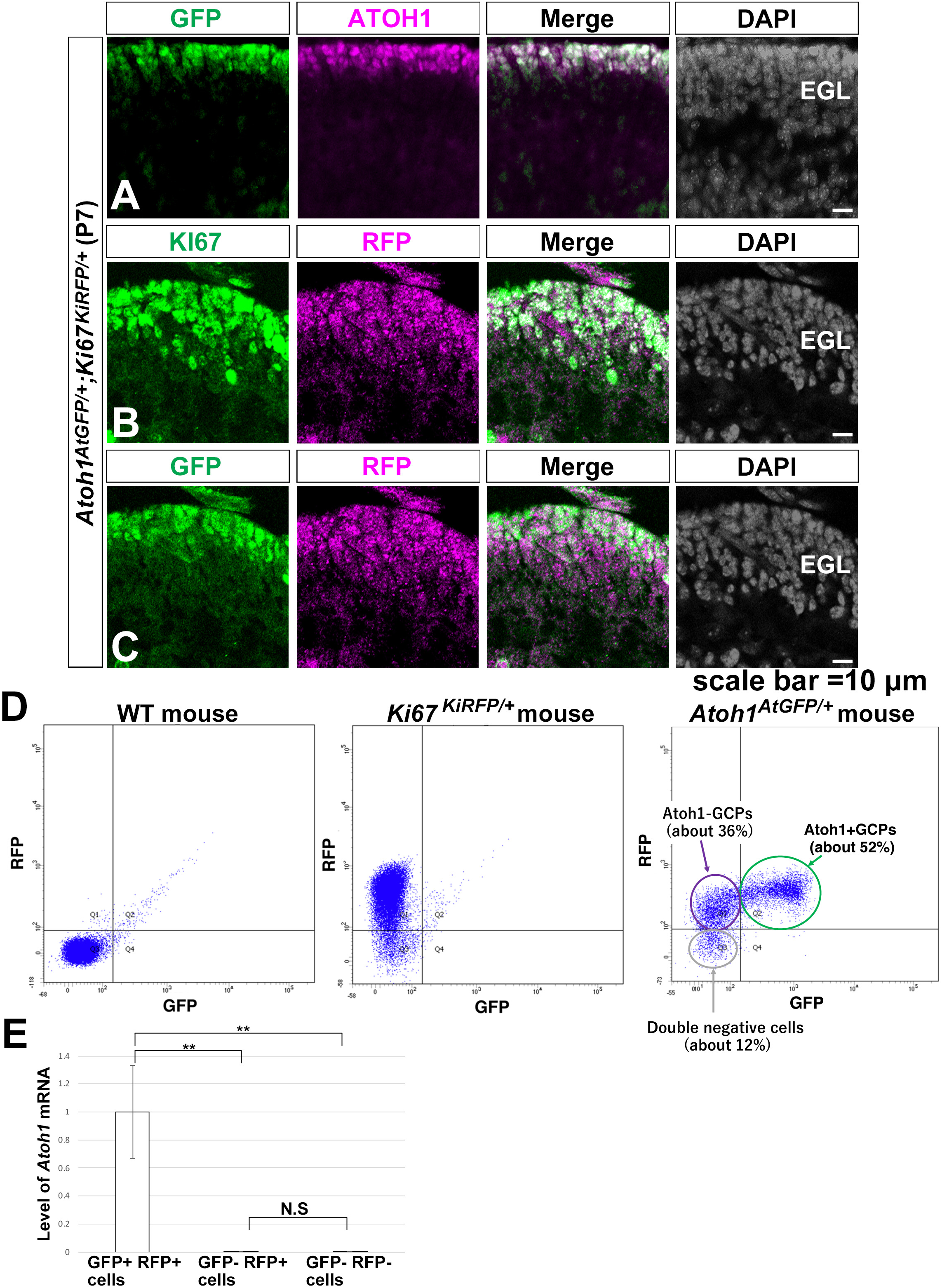
Isolation of distinct GCPs using *Atoh1^AtGFP/+^; Ki67^KiRFP /+^* reporter mice. **A.** Validation of the Atoh1-GFP reporter in the P7 cerebellum of an *Atoh1^AtGFP/+^; Ki67^KiRFP /+^* mouse. Representative images show co-localization of the GFP signal (green) with immunostaining for the endogenous ATOH1 protein (magenta). **B.** Validation of the Ki67-RFP reporter in the P7 cerebellum of an *Atoh1^AtGFP/+^; Ki67^KiRFP /+^*mouse. Representative images show co-localization of the RFP signal (magenta) with immunostaining for the endogenous KI67 protein (green). **C.** Combined expression patterns of ATOH1-GFP (green) and KI67-RFP (magenta) in the P7 cerebellum. The images show that while both reporters are expressed predominantly in the outer external granular layer (oEGL), ATOH1-GFP expression is more tightly restricted to the outermost cell layer compared to the broader expression of KI67-RFP. **D.** Representative FACS plots demonstrating the gating strategy for isolating distinct granule cell lineage populations from purified GCPs of Atoh1^AtGFP/+^; Ki67^KiRFP /+^ mice. Wild-type (WT) GCPs are used for negative control gating. **E.** Relative expression levels of Atoh1 transcripts, estimated by qRT-PCR, in FACS-sorted GCP populations purified from P6 cerebella. Atoh1 expression is exclusively detected in the GFP+RFP+ fraction, confirming the specificity of this highly proliferative population.

**Figure S4.**
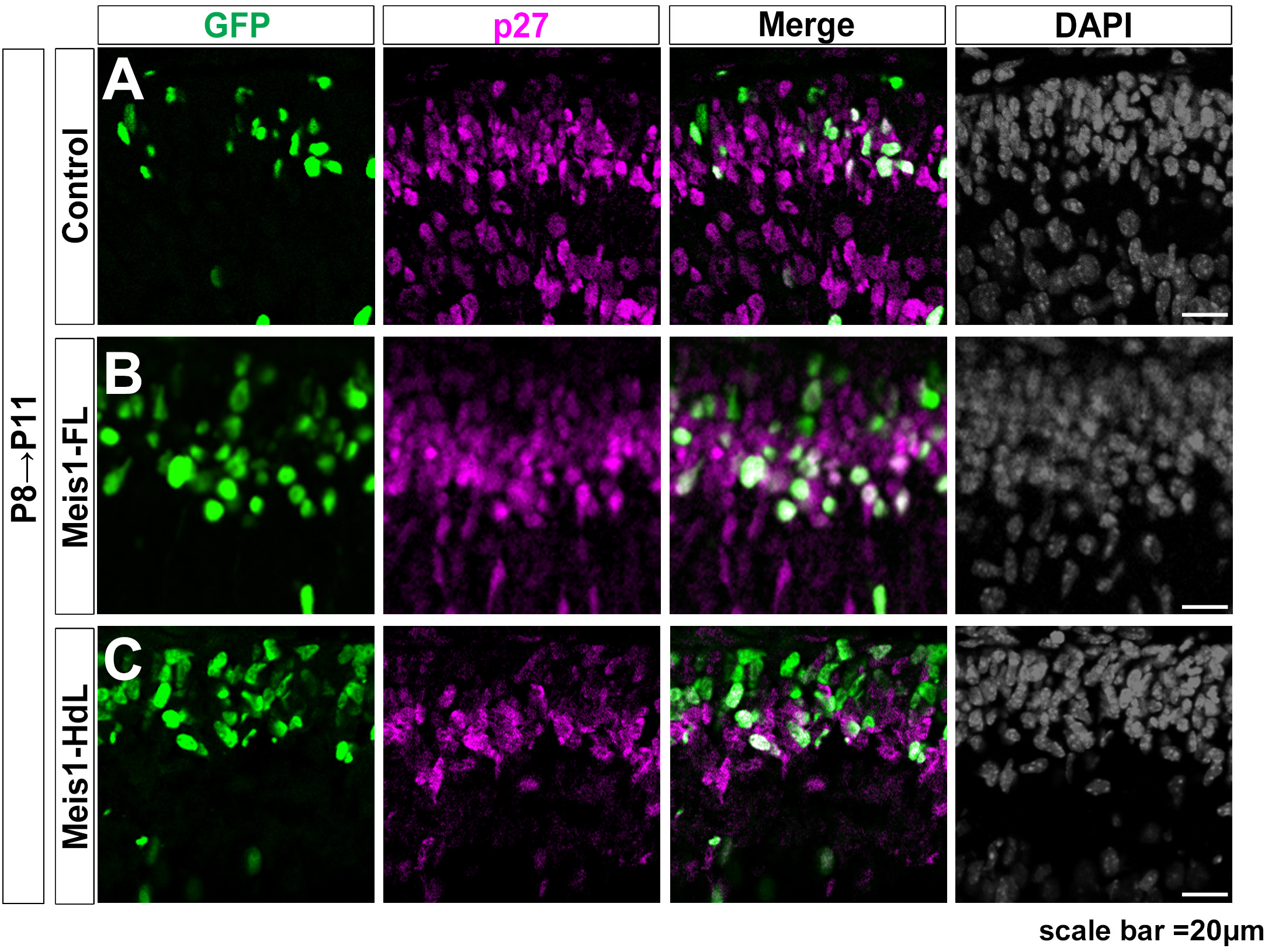
*Meis1* isoforms differentially regulate p27+ cell proportions *in vivo*. **A–C.** Representative immunofluorescence images of P11 cerebella following *in vivo* electroporation at P8. Cerebellar sections were immunostained for p27 (magenta). Electroporated cells, identified by co-electroporated H3.1-EGFP (green), were transfected with a control vector (A), MEIS1-FL (B), or MEIS1-HdL (C).

**Figure S5.**
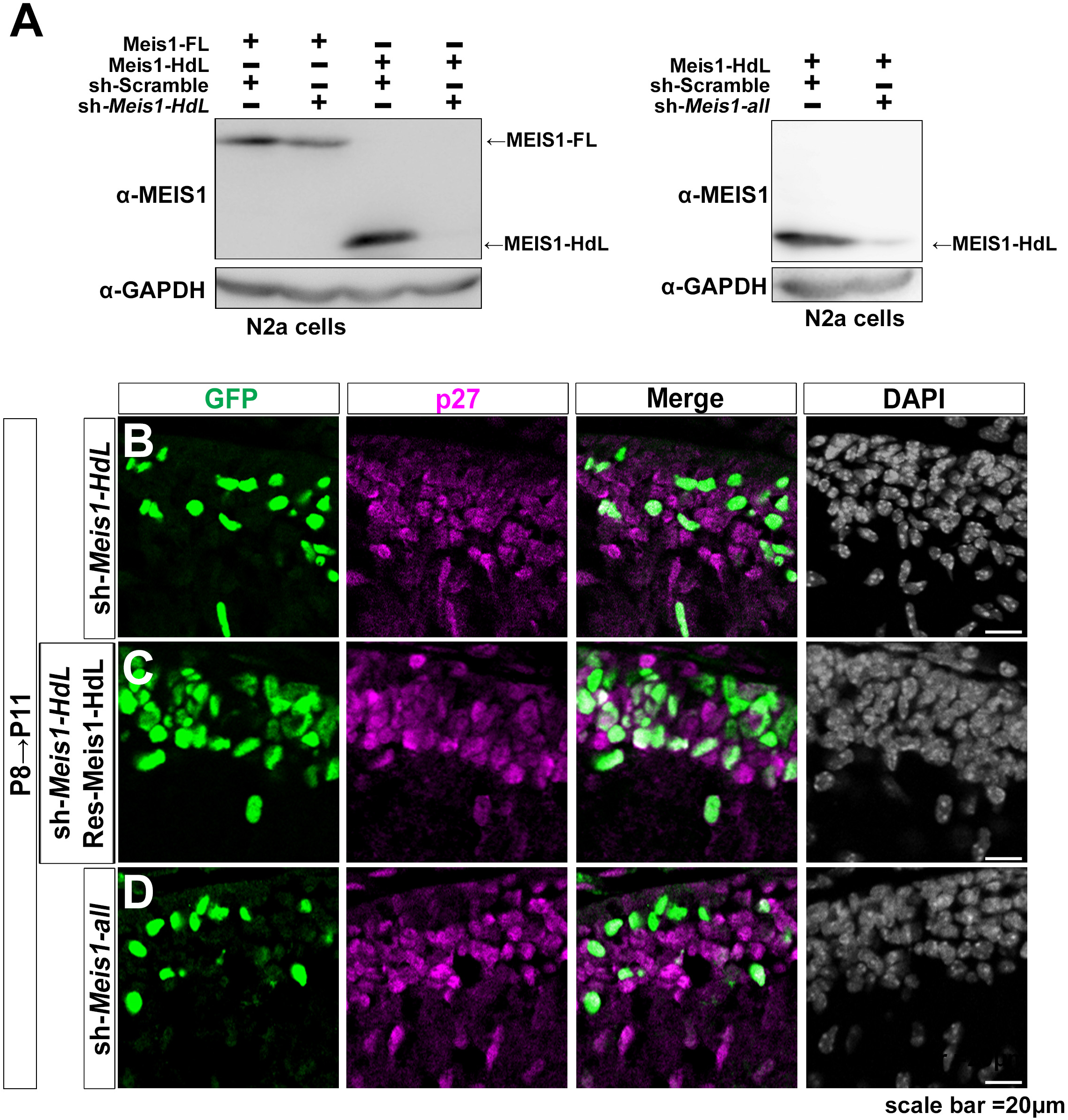
*Meis1-HdL* knockdown promotes GCP differentiation. **A.** Immunoblot analysis validating the specificity and efficacy of the indicated shRNAs in N2a cell lysates. Left panel: Specificity test for sh-*Meis1-HdL*. The blot shows that sh-*Meis1-HdL* targets *Meis1-HdL* for knockdown but does not affect *Meis1-FL*. Right panel: Efficacy test for sh-*Meis1-all*. The blot confirms that sh-*Meis1-all* effectively knocks down *Meis1-HdL*. **B–D.** Representative immunofluorescence images of P11 cerebella following *in vivo* electroporation at P8. Cerebellar sections were immunostained for p27 (magenta). Electroporated cells, identified by co-electroporated H3.1-EGFP (green), were transfected with (B) sh-*Meis1-HdL*, (C) sh-*Meis1-HdL* plus a knockdown-resistant rescue construct (Res-*Meis1-HdL*), or (D) sh-*Meis1-all*.

**Figure S6.**
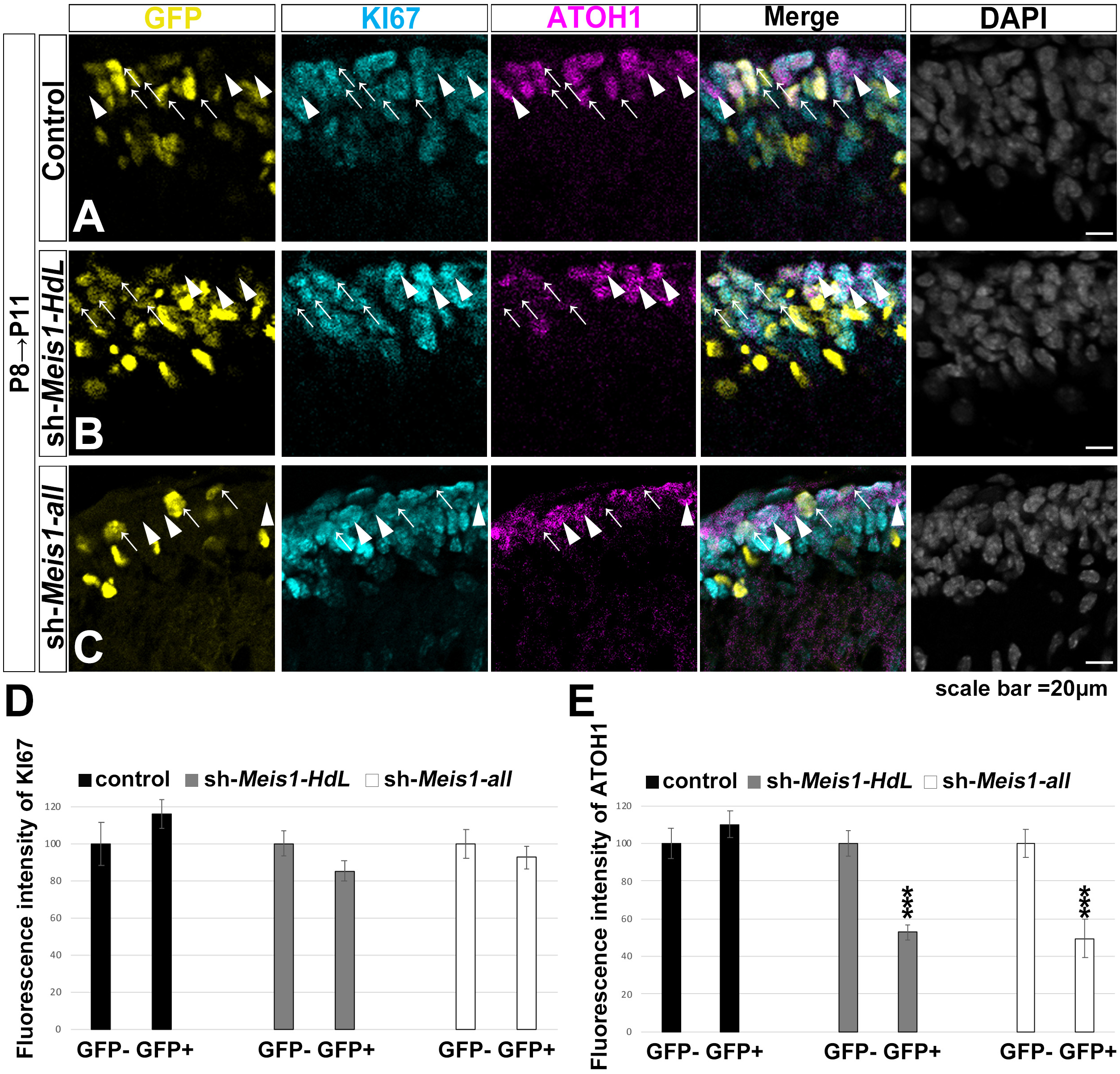
*Meis1* knockdown reduces ATOH1 protein levels in GCPs. **A–C.** Representative immunofluorescence images of P11 cerebella following *in vivo* electroporation at P8. Sections were immunostained for ATOH1 (magenta) and KI67 (cyan). Electroporated cell nuclei are identified by H3.1-EGFP (yellow). The images show cells transfected with a control vector (A), sh-*Meis1-HdL* (B), or sh-*Meis1-all* (C). Arrows indicate electroporated (GFP+) cells, and arrowheads indicate surrounding non-electroporated (GFP-) cells. **D.** Quantification of KI67 fluorescence intensity in electroporated (GFP+) and neighboring non-electroporated (GFP-) cells from the outer EGL, as shown in (A–C). The analysis indicates no significant change in KI67 levels following *Meis1* knockdown. **E.** Quantification of ATOH1 fluorescence intensity in electroporated (GFP+) and neighboring non-electroporated (GFP-) cells from the outer EGL, as shown in (A–C). The results show a significant reduction in ATOH1 levels in cells with *Meis1* knockdown.

**Figure S7.**
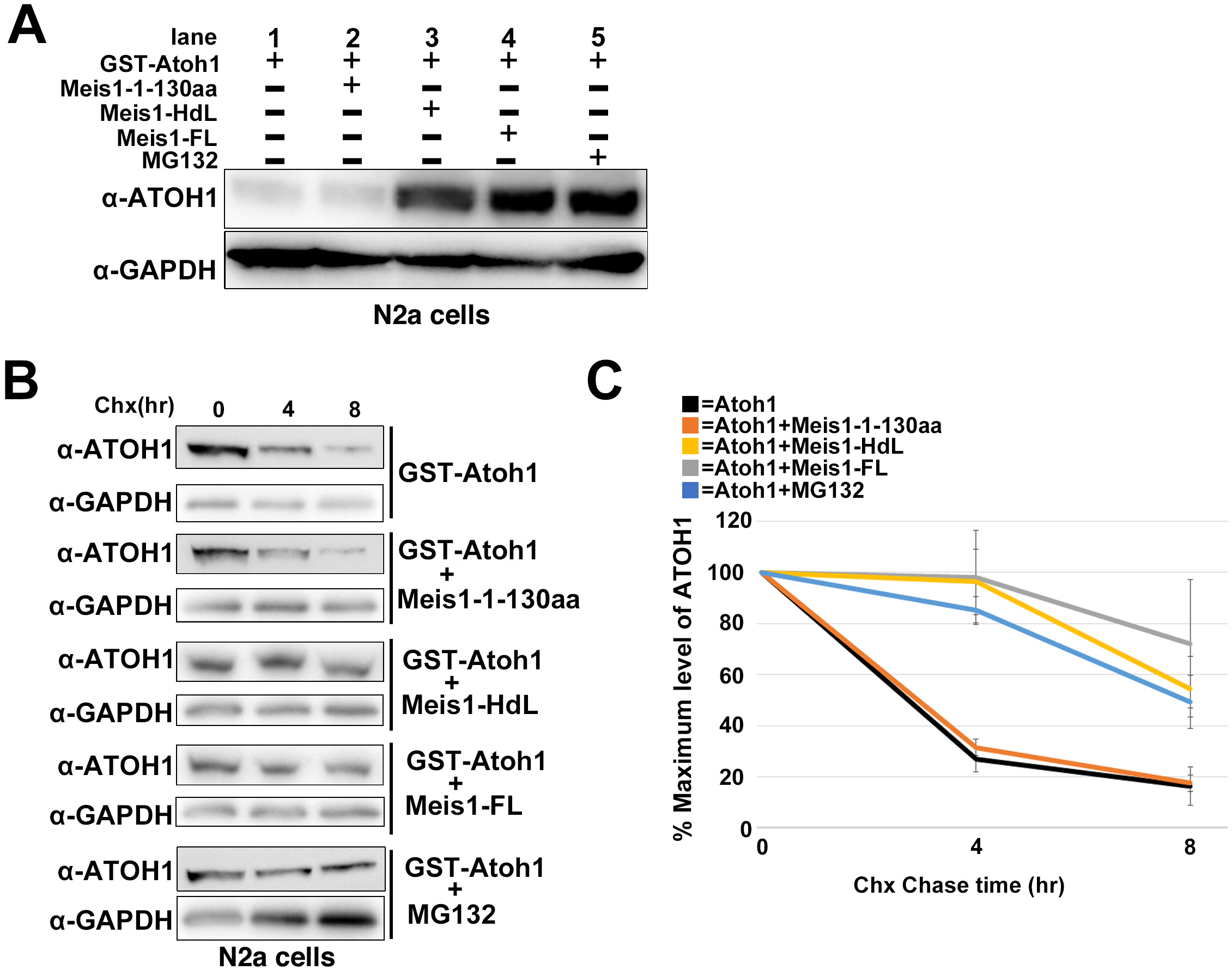
MEIS1 isoforms stabilize ATOH1 levels in a non-transcriptional manner. **A.** Immunoblotting analysis of N2a cell lysates. Cells were transfected with GST-ATOH1 and either no additional plasmid (control), Meis1-1–130aa fragment, Meis1-HdL, or Meis1-FL expression plasmids. Where indicated, cells were treated with MG132 for 6 hours prior to lysis. ATOH1 protein levels were determined using an anti-ATOH1 antibody, with GAPDH as a loading control. **B.** Immunoblotting analysis of N2a cell lysates following CHX chase assay. Cells were transfected with GST-ATOH1 and either Meis1-1–130aa fragment, Meis1-HdL, or Meis1-FL expression plasmids. Where indicated, cells were pre-treated with MG132 for 6 hours before CHX administration. Lysates were collected at 0, 4, and 8 hours after CHX treatment. ATOH1 protein levels were determined using an anti-ATOH1 antibody, with GAPDH as a loading control. **C.** Quantification of ATOH1 protein levels from (B), normalized to GAPDH.

**Figure S8.**
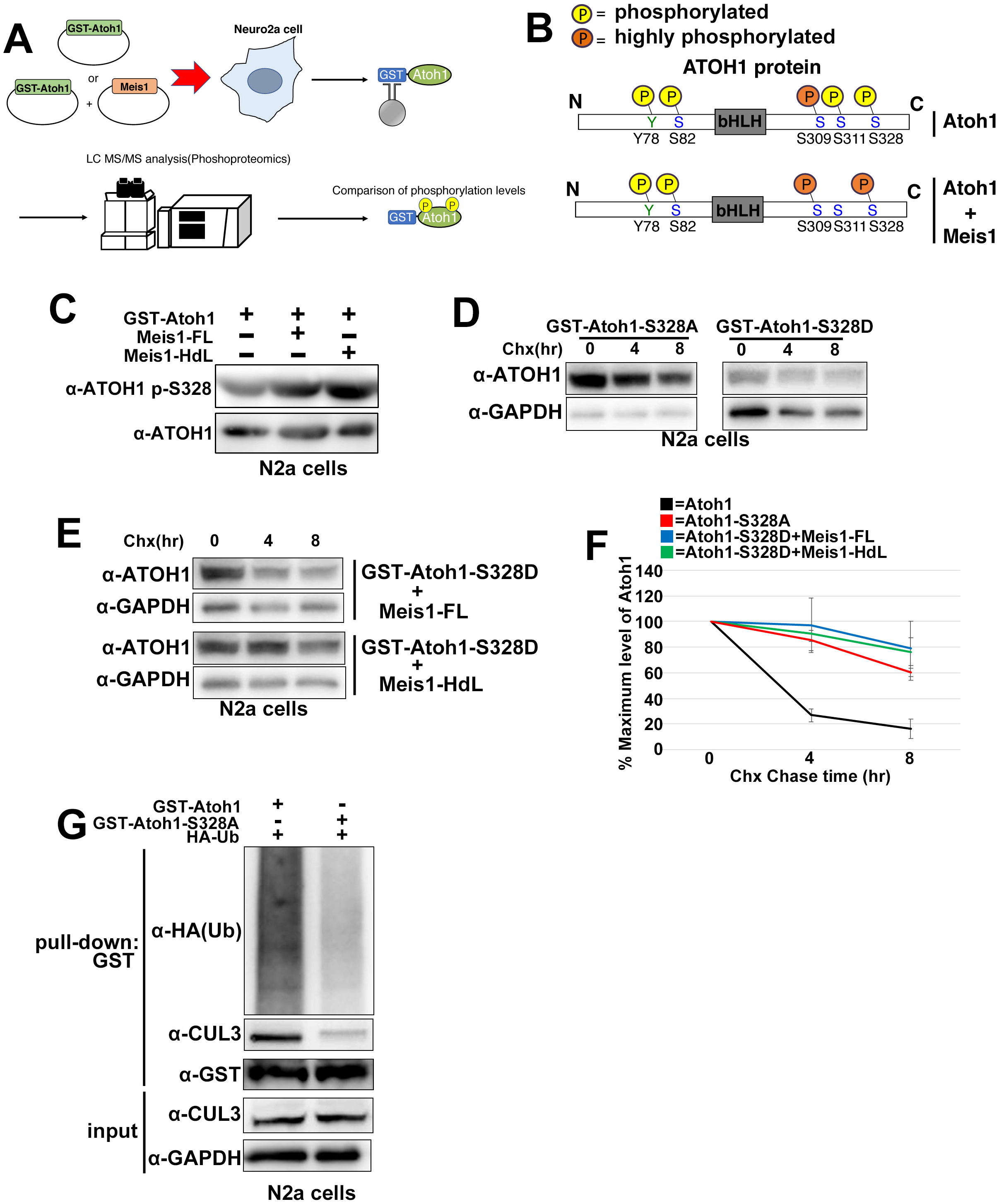
CUL3 recognizes and degrades S328 phosphorylated ATOH1, and MEIS1 inhibits the degradation of phosphorylated ATOH1. **A.** Schematic illustration of the proteomic analysis strategy used to identify phosphorylation sites of ATOH1 in N2a cells in the presence or absence of MEIS1. **B.** Schematic diagram summarizing ATOH1 phosphorylation sites identified by LC–MS/MS analysis. GST-ATOH1 was purified from N2a cells expressing it either alone or with MEIS1. The analysis identified five phosphorylation sites and revealed a marked increase in phosphorylation at the S328 residue in the presence of MEIS1. See Supplementary File for list of ATOH1 phosphorylation sites for complete data. **C.** Immunoblotting analysis of N2a cell lysates. Cells were transfected with GST-ATOH1 alone or in combination with MEIS1-FL or MEIS1-HdL. Samples were loaded to contain equal amounts of total ATOH1. ATOH1-S328 phosphorylation levels were determined by Western blotting with an anti-ATOH1 p-S328 antibody. **D.** Immunoblotting analysis of N2a cell lysates following CHX chase assay. Cells were transfected with GST-ATOH1-S328A (non-phosphorylated form) or GST-ATOH1-S328D (phosphomimic form) plasmids and harvested at 0, 4, and 8 hours after treatment with CHX. ATOH1 protein levels were detected with an anti-ATOH1 antibody, with GAPDH as a loading control. **E.** Immunoblotting analysis of N2a cell lysates following CHX chase assay. Cells were transfected with GST-ATOH1-S328D in combination with MEIS1-FL or MEIS1-HdL plasmids and harvested at 0, 4, and 8 hours after treatment with CHX. ATOH1 protein levels were detected with an anti-ATOH1 antibody, with GAPDH as a loading control. **F.** Quantification of ATOH1 protein levels from (D) and (E), normalized to GAPDH. **G.** Immunoblotting analysis of GST pull-down fractions from N2a cells co-transfected with HA-Ub and either GST-ATOH1 (wild-type) or GST-ATOH1-S328A. Polyubiquitinated ATOH1 and total GST-ATOH1 levels were determined by immunoblotting with anti-HA and anti-GST antibodies, respectively. The interaction of CUL3 with GST-ATOH1 was assessed by immunoblotting with an anti-CUL3 antibody.

**Figure S9.**
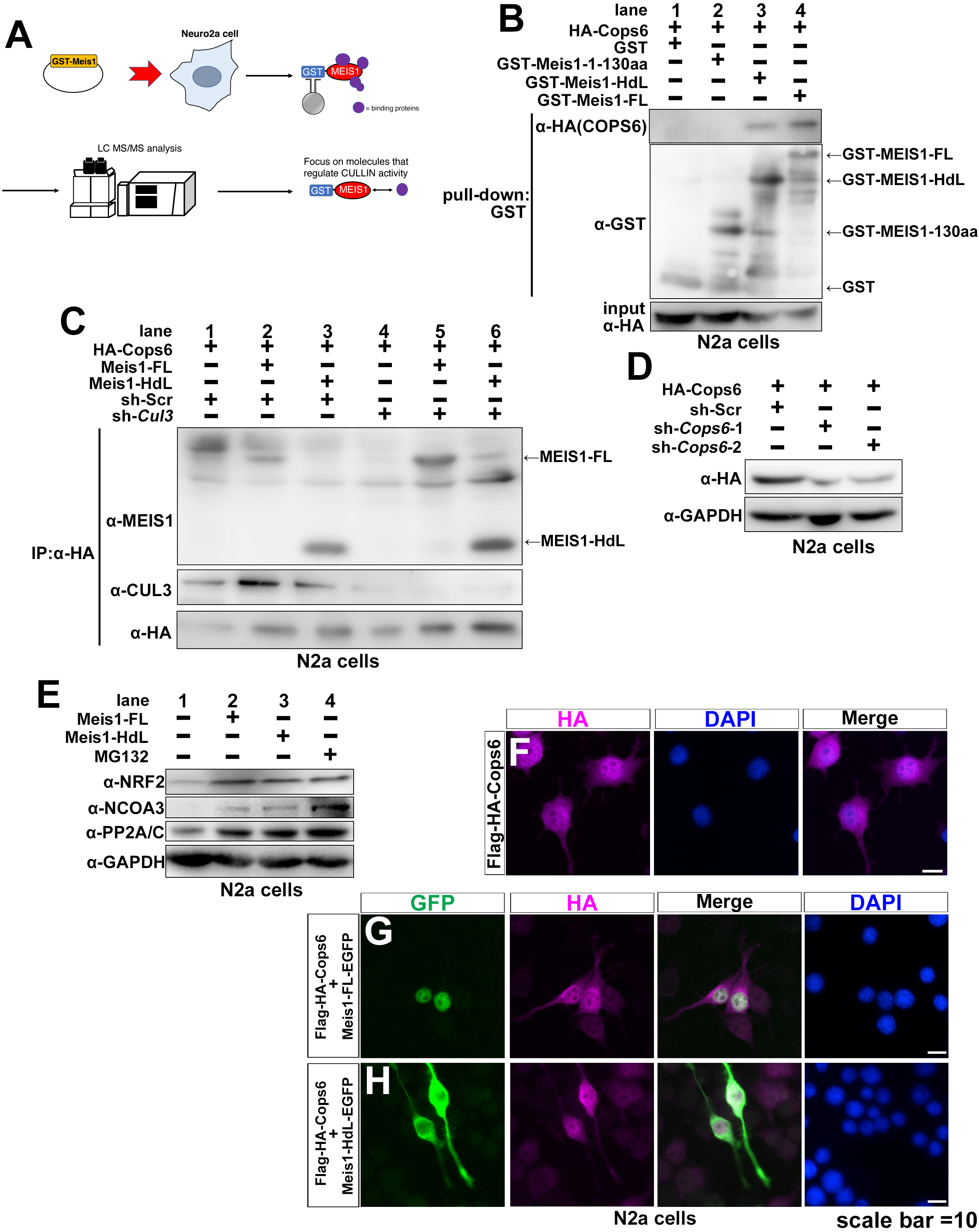
MEIS1 regulates not only ATOH1 but also other CUL3 target proteins. **A.** Schematic illustrating the proteomic analysis strategy used to identify MEIS1-binding molecules. **B.** Immunoblotting analysis of GST pull-down fractions from N2a cells transfected with GST, GST-MEIS1-1–130aa fragment, GST-MEIS1-HdL, or GST-MEIS1-FL. All cells were also co-transfected with HA-COPS6. The blot, probed for HA, shows that COPS6 binds to MEIS1-FL and MEIS1-HdL, but not to the N-terminal (1–130aa) fragment or the GST control. **C.** Co-immunoprecipitation assay demonstrating that the MEIS1–COPS6 interaction is independent of CUL3. HA-COPS6 was immunoprecipitated from N2a cells co-expressing a MEIS1 isoform along with either a control scramble shRNA or an shRNA targeting *Cul3*. The immunoblots show that the amount of co-precipitated MEIS1 is unaffected by the knockdown of *Cul3*. **D.** Immunoblotting analysis of N2a cell lysates. Cells were co-transfected with HA-COPS6 and either a control scramble shRNA or one of two shRNAs targeting Cops6 (sh-*Cops6-1* and sh-*Cops6-2*). The blot was probed with an anti-HA antibody and shows that both shRNAs effectively knock down HA-COPS6 expression compared to the control. **E.** Immunoblot analysis showing that MEIS1 isoforms stabilize known CUL3 target proteins. N2a cells were transfected with the indicated MEIS1 constructs. Treatment with the proteasome inhibitor MG132 was used as a positive control. The blots show that overexpression of either MEIS1-FL or MEIS1-HdL increases the protein levels of CUL3 targets (NRF2, NCOA3, PP2A/C), similar to the effect of MG132. **F–H.** Representative immunofluorescence images showing the subcellular localization of HA-COPS6 in N2a cells. Cells were transfected with HA-COPS6 alone (F), or co-transfected with HA-COPS6 and MEIS1-FL-EGFP (G), or MEIS1-HdL-EGFP (H). Cells were immunostained for HA (magenta), GFP (green, to visualize MEIS1-EGFP fusion proteins), and DAPI (blue, for nuclei).

## Reference

1. Wright, C.J., Smith, C.W.J., and Jiggins, C.D. (2022). Alternative splicing as a source of phenotypic diversity. Nat Rev Genet 23, 697–710. 10.1038/s41576-022-00514-4.

2. Marasco, L.E., and Kornblihtt, A.R. (2023). The physiology of alternative splicing. Nat Rev Mol Cell Biol 24, 242–254. 10.1038/s41580-022-00545-z.

3. LaForce, G.R., Philippidou, P., and Schaffer, A.E. (2023). mRNA isoform balance in neuronal development and disease. Wiley Interdiscip Rev RNA 14, e1762. 10.1002/wrna.1762.

4. Patowary, A., Zhang, P., Jops, C., Vuong, C.K., Ge, X., Hou, K., Kim, M., Gong, N., Margolis, M., Vo, D., et al. (2024). Developmental isoform diversity in the human neocortex informs neuropsychiatric risk mechanisms. Science 384, eadh7688. 10.1126/science.adh7688.

5. Joglekar, A., Hu, W., Zhang, B., Narykov, O., Diekhans, M., Marrocco, J., Balacco, J., Ndhlovu, L.C., Milner, T.A., Fedrigo, O., et al. (2024). Single-cell long-read sequencing-based mapping reveals specialized splicing patterns in developing and adult mouse and human brain. Nat Neurosci 27, 1051–1063. 10.1038/s41593-024-01616-4.

6. De Paoli-Iseppi, R., Gleeson, J., and Clark, M.B. (2021). Isoform Age - Splice Isoform Profiling Using Long-Read Technologies. Front Mol Biosci 8, 711733. 10.3389/fmolb.2021.711733.

7. Monzo, C., Liu, T., and Conesa, A. (2025). Transcriptomics in the era of long-read sequencing. Nat Rev Genet. 10.1038/s41576-025-00828-z.

8. Miyashita, S., Owa, T., Seto, Y., Yamashita, M., Aida, S., Sone, M., Ichijo, K., Nishioka, T., Kaibuchi, K., Kawaguchi, Y., et al. (2021). Cyclin D1 controls development of cerebellar granule cell progenitors through phosphorylation and stabilization of ATOH1. EMBO J 40, e105712. 10.15252/embj.2020105712.

9. Miyashita, S., and Hoshino, M. (2022). Transit Amplifying Progenitors in the Cerebellum: Similarities to and Differences from Transit Amplifying Cells in Other Brain Regions and between Species. Cells 11. 10.3390/cells11040726.

10. Forget, A., Bihannic, L., Cigna, S.M., Lefevre, C., Remke, M., Barnat, M., Dodier, S., Shirvani, H., Mercier, A., Mensah, A., et al. (2014). Shh signaling protects Atoh1 from degradation mediated by the E3 ubiquitin ligase Huwe1 in neural precursors. Dev Cell 29, 649–661. 10.1016/j.devcel.2014.05.014.

11. Zhao, H., Ayrault, O., Zindy, F., Kim, J.H., and Roussel, M.F. (2008). Post-transcriptional down-regulation of Atoh1/Math1 by bone morphogenic proteins suppresses medulloblastoma development. Genes Dev 22, 722–727. 10.1101/gad.1636408.

12. Schulte, D., and Geerts, D. (2019). MEIS transcription factors in development and disease. Development 146. 10.1242/dev.174706.

13. Heine, P., Dohle, E., Bumsted-O’Brien, K., Engelkamp, D., and Schulte, D. (2008). Evidence for an evolutionary conserved role of homothorax/Meis1/2 during vertebrate retina development. Development 135, 805–811. 10.1242/dev.012088.

14. Tucker, E.S., Lehtinen, M.K., Maynard, T., Zirlinger, M., Dulac, C., Rawson, N., Pevny, L., and Lamantia, A.S. (2010). Proliferative and transcriptional identity of distinct classes of neural precursors in the mammalian olfactory epithelium. Development 137, 2471–2481. 10.1242/dev.049718.

15. Azcoitia, V., Aracil, M., Martinez, A.C., and Torres, M. (2005). The homeodomain protein Meis1 is essential for definitive hematopoiesis and vascular patterning in the mouse embryo. Dev Biol 280, 307–320. 10.1016/j.ydbio.2005.01.004.

16. Owa, T., Taya, S., Miyashita, S., Yamashita, M., Adachi, T., Yamada, K., Yokoyama, M., Aida, S., Nishioka, T., Inoue, Y.U., et al. (2018). Meis1 Coordinates Cerebellar Granule Cell Development by Regulating Pax6 Transcription, BMP Signaling and Atoh1 Degradation. J Neurosci 38, 1277–1294. 10.1523/JNEUROSCI.1545-17.2017.

17. Wang, Y., Zhao, Y., Bollas, A., Wang, Y., and Au, K.F. (2021). Nanopore sequencing technology, bioinformatics and applications. Nat Biotechnol 39, 1348–1365. 10.1038/s41587-021-01108-x.

18. Engelkamp, D., Rashbass, P., Seawright, A., and van Heyningen, V. (1999). Role of Pax6 in development of the cerebellar system. Development 126, 3585–3596. 10.1242/dev.126.16.3585.

19. Swanson, D.J., and Goldowitz, D. (2011). Experimental Sey mouse chimeras reveal the developmental deficiencies of Pax6-null granule cells in the postnatal cerebellum. Dev Biol 351, 1–12. 10.1016/j.ydbio.2010.11.018.

20. Steele-Perkins, G., Plachez, C., Butz, K.G., Yang, G., Bachurski, C.J., Kinsman, S.L., Litwack, E.D., Richards, L.J., and Gronostajski, R.M. (2005). The transcription factor gene Nfib is essential for both lung maturation and brain development. Mol Cell Biol 25, 685–698. 10.1128/MCB.25.2.685-698.2005.

21. Crist, R.C., Roth, J.J., Waldman, S.A., and Buchberg, A.M. (2011). A conserved tissue-specific homeodomain-less isoform of MEIS1 is downregulated in colorectal cancer. PLoS One 6, e23665. 10.1371/journal.pone.0023665.

22. Xie, Q., Ung, D., Khafizov, K., Fiser, A., and Cvekl, A. (2014). Gene regulation by PAX6: structural-functional correlations of missense mutants and transcriptional control of Trpm3/miR-204. Mol Vis 20, 270–282.

23. Azuma, N., Tadokoro, K., Asaka, A., Yamada, M., Yamaguchi, Y., Handa, H., Matsushima, S., Watanabe, T., Kohsaka, S., Kida, Y., et al. (2005). The Pax6 isoform bearing an alternative spliced exon promotes the development of the neural retinal structure. Hum Mol Genet 14, 735–745. 10.1093/hmg/ddi069.

24. Liu, Y., Bernard, H.U., and Apt, D. (1997). NFI-B3, a novel transcriptional repressor of the nuclear factor I family, is generated by alternative RNA processing. J Biol Chem 272, 10739–10745. 10.1074/jbc.272.16.10739.

25. Chen, L., Kostadima, M., Martens, J.H.A., Canu, G., Garcia, S.P., Turro, E., Downes, K., Macaulay, I.C., Bielczyk-Maczynska, E., Coe, S., et al. (2014). Transcriptional diversity during lineage commitment of human blood progenitors. Science 345, 1251033. 10.1126/science.1251033.

26. Rose, M.F., Ren, J., Ahmad, K.A., Chao, H.T., Klisch, T.J., Flora, A., Greer, J.J., and Zoghbi, H.Y. (2009). Math1 is essential for the development of hindbrain neurons critical for perinatal breathing. Neuron 64, 341–354. 10.1016/j.neuron.2009.10.023.

27. Basak, O., van de Born, M., Korving, J., Beumer, J., van der Elst, S., van Es, J.H., and Clevers, H. (2014). Mapping early fate determination in Lgr5+ crypt stem cells using a novel Ki67-RFP allele. EMBO J 33, 2057–2068. 10.15252/embj.201488017.

28. Flora, A., Klisch, T.J., Schuster, G., and Zoghbi, H.Y. (2009). Deletion of Atoh1 disrupts Sonic Hedgehog signaling in the developing cerebellum and prevents medulloblastoma. Science 326, 1424–1427. 10.1126/science.1181453.

29. Helms, A.W., Gowan, K., Abney, A., Savage, T., and Johnson, J.E. (2001). Overexpression of MATH1 disrupts the coordination of neural differentiation in cerebellum development. Mol Cell Neurosci 17, 671–682. 10.1006/mcne.2000.0969.

30. Chang, C.H., Zanini, M., Shirvani, H., Cheng, J.S., Yu, H., Feng, C.H., Mercier, A.L., Hung, S.Y., Forget, A., Wang, C.H., et al. (2019). Atoh1 Controls Primary Cilia Formation to Allow for SHH-Triggered Granule Neuron Progenitor Proliferation. Dev Cell 48, 184–199 e185. 10.1016/j.devcel.2018.12.017.

31. Wu, C., Macleod, I., and Su, A.I. (2013). BioGPS and MyGene.info: organizing online, gene-centric information. Nucleic Acids Res 41, D561–565. 10.1093/nar/gks1114.

32. Cheng, Y.F., Tong, M., and Edge, A.S. (2016). Destabilization of Atoh1 by E3 Ubiquitin Ligase Huwe1 and Casein Kinase 1 Is Essential for Normal Sensory Hair Cell Development. J Biol Chem 291, 21096–21109. 10.1074/jbc.M116.722124.

33. Aragaki, M., Tsuchiya, K., Okamoto, R., Yoshioka, S., Nakamura, T., Sakamoto, N., Kanai, T., and Watanabe, M. (2008). Proteasomal degradation of Atoh1 by aberrant Wnt signaling maintains the undifferentiated state of colon cancer. Biochem Biophys Res Commun 368, 923–929. 10.1016/j.bbrc.2008.02.011.

34. Baek, K., Scott, D.C., and Schulman, B.A. (2021). NEDD8 and ubiquitin ligation by cullin-RING E3 ligases. Curr Opin Struct Biol 67, 101–109. 10.1016/j.sbi.2020.10.007.

35. Chua, Y.S., Boh, B.K., Ponyeam, W., and Hagen, T. (2011). Regulation of cullin RING E3 ubiquitin ligases by CAND1 in vivo. PLoS One 6, e16071. 10.1371/journal.pone.0016071.

36. Petroski, M.D., and Deshaies, R.J. (2005). Function and regulation of cullin-RING ubiquitin ligases. Nat Rev Mol Cell Biol 6, 9–20. 10.1038/nrm1547.

37. Tsuchiya, K., Nakamura, T., Okamoto, R., Kanai, T., and Watanabe, M. (2007). Reciprocal targeting of Hath1 and beta-catenin by Wnt glycogen synthase kinase 3beta in human colon cancer. Gastroenterology 132, 208–220. 10.1053/j.gastro.2006.10.031.

38. Jaramillo, M.C., and Zhang, D.D. (2013). The emerging role of the Nrf2-Keap1 signaling pathway in cancer. Genes Dev 27, 2179–2191. 10.1101/gad.225680.113.

39. Ferry, C., Gaouar, S., Fischer, B., Boeglin, M., Paul, N., Samarut, E., Piskunov, A., Pankotai-Bodo, G., Brino, L., and Rochette-Egly, C. (2011). Cullin 3 mediates SRC-3 ubiquitination and degradation to control the retinoic acid response. Proc Natl Acad Sci U S A 108, 20603–20608. 10.1073/pnas.1102572108.

40. Xu, J., Zhou, J.Y., Xu, Z., Kho, D.H., Zhuang, Z., Raz, A., and Wu, G.S. (2014). The role of Cullin3-mediated ubiquitination of the catalytic subunit of PP2A in TRAIL signaling. Cell Cycle 13, 3750–3758. 10.4161/15384101.2014.965068.

41. Cullinan, S.B., Gordan, J.D., Jin, J., Harper, J.W., and Diehl, J.A. (2004). The Keap1-BTB protein is an adaptor that bridges Nrf2 to a Cul3-based E3 ligase: oxidative stress sensing by a Cul3-Keap1 ligase. Mol Cell Biol 24, 8477–8486. 10.1128/MCB.24.19.8477-8486.2004.

42. Schulze-Niemand, E., and Naumann, M. (2023). The COP9 signalosome: A versatile regulatory hub of Cullin-RING ligases. Trends Biochem Sci 48, 82–95. 10.1016/j.tibs.2022.08.003.

43. Rao, F., Xu, J., Khan, A.B., Gadalla, M.M., Cha, J.Y., Xu, R., Tyagi, R., Dang, Y., Chakraborty, A., and Snyder, S.H. (2014). Inositol hexakisphosphate kinase-1 mediates assembly/disassembly of the CRL4-signalosome complex to regulate DNA repair and cell death. Proc Natl Acad Sci U S A 111, 16005–16010. 10.1073/pnas.1417900111.

44. Scherer, P.C., Ding, Y., Liu, Z., Xu, J., Mao, H., Barrow, J.C., Wei, N., Zheng, N., Snyder, S.H., and Rao, F. (2016). Inositol hexakisphosphate (IP6) generated by IP5K mediates cullin-COP9 signalosome interactions and CRL function. Proc Natl Acad Sci U S A 113, 3503–3508. 10.1073/pnas.1525580113.

45. Lee, M.H., Zhao, R., Phan, L., and Yeung, S.C. (2011). Roles of COP9 signalosome in cancer. Cell Cycle 10, 3057–3066. 10.4161/cc.10.18.17320.

46. Liu, J., Qin, Y.Z., Yang, S., Wang, Y., Chang, Y.J., Zhao, T., Jiang, Q., and Huang, X.J. (2017). Meis1 is critical to the maintenance of human acute myeloid leukemia cells independent of MLL rearrangements. Ann Hematol 96, 567–574. 10.1007/s00277-016-2913-6.

47. Liu, G., Claret, F.X., Zhou, F., and Pan, Y. (2018). Jab1/COPS5 as a Novel Biomarker for Diagnosis, Prognosis, Therapy Prediction and Therapeutic Tools for Human Cancer. Front Pharmacol 9, 135. 10.3389/fphar.2018.00135.

48. Mori, M., Yoneda-Kato, N., Yoshida, A., and Kato, J.Y. (2008). Stable form of JAB1 enhances proliferation and maintenance of hematopoietic progenitors. J Biol Chem 283, 29011–29021. 10.1074/jbc.M804539200.

49. Thorsteinsdottir, U., Kroon, E., Jerome, L., Blasi, F., and Sauvageau, G. (2001). Defining roles for HOX and MEIS1 genes in induction of acute myeloid leukemia. Mol Cell Biol 21, 224–234. 10.1128/MCB.21.1.224-234.2001.

50. Wang, J.X., and Furukawa, H. (2019). Dissecting diverse functions of NMDA receptors by structural biology. Curr Opin Struct Biol 54, 34–42. 10.1016/j.sbi.2018.12.009.

51. Li, H., Rajani, V., Sengar, A.S., and Salter, M.W. (2024). Src dependency of the regulation of LTP by alternative splicing of GRIN1 exon 5. Philos Trans R Soc Lond B Biol Sci 379, 20230236. 10.1098/rstb.2023.0236.

52. Zhou, L., and Duan, J. (2018). The C-terminus of NMDAR GluN1-1a Subunit Translocates to Nucleus and Regulates Synaptic Function. Front Cell Neurosci 12, 334. 10.3389/fncel.2018.00334.

53. Llansola, M., Sanchez-Perez, A., Cauli, O., and Felipo, V. (2005). Modulation of NMDA receptors in the cerebellum. 1. Properties of the NMDA receptor that modulate its function. Cerebellum 4, 154–161. 10.1080/14734220510007996.

54. Adachi, T., Miyashita, S., Yamashita, M., Shimoda, M., Okonechnikov, K., Chavez, L., Kool, M., Pfister, S.M., Inoue, T., Kawauchi, D., and Hoshino, M. (2021). Notch Signaling between Cerebellar Granule Cell Progenitors. eNeuro 8. 10.1523/ENEURO.0468-20.2021.

55. Kutscher, L.M., Okonechnikov, K., Batora, N.V., Clark, J., Silva, P.B.G., Vouri, M., van Rijn, S., Sieber, L., Statz, B., Gearhart, M.D., et al. (2020). Functional loss of a noncanonical BCOR-PRC1.1 complex accelerates SHH-driven medulloblastoma formation. Genes Dev 34, 1161–1176. 10.1101/gad.337584.120.

56. Prjibelski, A.D., Mikheenko, A., Joglekar, A., Smetanin, A., Jarroux, J., Lapidus, A.L., and Tilgner, H.U. (2023). Accurate isoform discovery with IsoQuant using long reads. Nat Biotechnol 41, 915–918. 10.1038/s41587-022-01565-y.

57. Li, H. (2021). New strategies to improve minimap2 alignment accuracy. Bioinformatics 37, 4572–4574. 10.1093/bioinformatics/btab705.

58. Pardo-Palacios, F.J., Arzalluz-Luque, A., Kondratova, L., Salguero, P., Mestre-Tomas, J., Amorin, R., Estevan-Morio, E., Liu, T., Nanni, A., McIntyre, L., et al. (2024). SQANTI3: curation of long-read transcriptomes for accurate identification of known and novel isoforms. Nat Methods 21, 793–797. 10.1038/s41592-024-02229-2.

59. Nagai, T., Nakamuta, S., Kuroda, K., Nakauchi, S., Nishioka, T., Takano, T., Zhang, X., Tsuboi, D., Funahashi, Y., Nakano, T., et al. (2016). Phosphoproteomics of the Dopamine Pathway Enables Discovery of Rap1 Activation as a Reward Signal In Vivo. Neuron 89, 550–565. 10.1016/j.neuron.2015.12.019.

60. Chen, E.Y., Tan, C.M., Kou, Y., Duan, Q., Wang, Z., Meirelles, G.V., Clark, N.R., and Ma’ayan, A. (2013). Enrichr: interactive and collaborative HTML5 gene list enrichment analysis tool. BMC Bioinformatics 14, 128. 10.1186/1471-2105-14-128.

61. Kuleshov, M.V., Jones, M.R., Rouillard, A.D., Fernandez, N.F., Duan, Q., Wang, Z., Koplev, S., Jenkins, S.L., Jagodnik, K.M., Lachmann, A., et al. (2016). Enrichr: a comprehensive gene set enrichment analysis web server 2016 update. Nucleic Acids Res 44, W90–97. 10.1093/nar/gkw377.

62. Xie, Z., Bailey, A., Kuleshov, M.V., Clarke, D.J.B., Evangelista, J.E., Jenkins, S.L., Lachmann, A., Wojciechowicz, M.L., Kropiwnicki, E., Jagodnik, K.M., et al. (2021). Gene Set Knowledge Discovery with Enrichr. Curr Protoc 1, e90. 10.1002/cpz1.90.

63. Gustavsson, E.K., Zhang, D., Reynolds, R.H., Garcia-Ruiz, S., and Ryten, M. (2022). ggtranscript: an R package for the visualization and interpretation of transcript isoforms using ggplot2. Bioinformatics 38, 3844–3846. 10.1093/bioinformatics/btac409.

64. Blum, M., Andreeva, A., Florentino, L.C., Chuguransky, S.R., Grego, T., Hobbs, E., Pinto, B.L., Orr, A., Paysan-Lafosse, T., Ponamareva, I., et al. (2025). InterPro: the protein sequence classification resource in 2025. Nucleic Acids Res 53, D444–D456. 10.1093/nar/gkae1082.

65. Brennan, P. (2018). drawProteins: a Bioconductor/R package for reproducible and programmatic generation of protein schematics. F1000Res 7, 1105. 10.12688/f1000research.14541.1.

